# Revealing the pervasive landscape of MGE-host interactions *in situ* with single-cell genomics

**DOI:** 10.64898/2025.12.20.690322

**Authors:** Ming Yan, Jill Banfield, Rohan Sachdeva

## Abstract

Mobile genetic elements (MGEs), including plasmids and viruses, drive microbial evolution and ecosystem dynamics, yet their distribution, host range, and functions remain poorly understood, especially among uncultivated lineages. Using ∼60,000 single-cell amplified genomes (SAGs) from host-associated and environmental microbiomes, we identified MGEs internal to or attached to individual cells, directly linking them to their hosts. Between 25% and 75% of cells contained at least one MGE, with gut-derived SAGs showing the highest MGE load. While most MGEs exhibited narrow, species-specific host ranges, a subset spanned multiple species and higher taxonomic ranks. We detected MGE clusters and complete, nearly identical genomes repeatedly associated with hosts across phyla, suggesting both genuinely broad host ranges and recurrent DNA entry events. We also identified microbial species acting as major donors or recipients in antibiotic resistance gene (ARG) exchange, and MGEs, including unclassified types that mediate extensive horizontal transfer of auxiliary functions. Together, these findings illuminate MGE–host associations and underscore that the genetic content of microbial cells is fundamentally linked to their MGEs in nature.

## Introduction

Metagenomics has transformed our understanding of microbial diversity^1–3^ and functional potential^4,5^. The massive scale of sequencing data has also enabled the discovery of diverse MGEs, including plasmids and viruses^6–8^, and offered insights into microbial-MGE ecology and evolutionary dynamics. Viral regulation of microbial metabolism significantly influences biogeochemical cycles^9,10^, has clinical relevance^11^, and also provides valuable genetic resources for biotechnological innovation^12^. Likewise, plasmid-mediated horizontal gene transfer (HGT) plays a pivotal role in microbial interactions, facilitating niche adaptation and promoting community-level metabolic cooperation, with broad ecological and clinical implications^13,14^. However, much remains to be learned about the auxiliary genetic repertoire of microbiomes that derives from MGEs because the association between specific MGEs and their hosts is unknown.

Various bioinformatic approaches have been developed to predict MGE-host associations within microbiomes, but suffer from limitations in recovery and accuracy of host assignments. Arguably, the highest fidelity approach involves matching CRISPR spacer sequences to MGEs as it assumes that possession of a spacer sequence is due to a past encounter with that MGE. However, this method is limited to microbes that possess CRISPR systems^15^ and often fails to uncover associations between coexisting viruses/phages and hosts because spacer targeting likely confers immunity. Moreover, recent findings suggest that such matches may reflect MGE entry without successful replication^16^. Machine learning–based host prediction tools based on phage–phage genomic similarity and phage–host signature concordance are limited, some achieve only genus-level resolution^17^, while others reach finer taxonomic levels but are restricted to a narrow range of pathogenic species^18,19^. In both cases, prediction accuracy declines significantly for novel or uncharacterized MGEs^17^. Consequently, the extent to which the apparent discrepancy between the broad host range of viruses identified in some metagenomes based on spacer matches^16,20^ and the predominantly strain-specific host associations observed in isolated viruses can be reconciled remains an open question.

High-resolution isolate sequencing, such as the analysis of over 1,300 *Escherichia coli* genomes, has shown that a typical *E. coli* genome contains at least one transposable element, a prophage, and a plasmid. At the pangenome level, MGEs account for more than one-third of the total gene content across *E. coli* phylogroups, highlighting their central role in shaping genomic diversity within this species^21^. Additionally, analysis of nearly 5,000 gut microbe isolate genomes revealed that 70% of the isolates, spanning over 70 genera, contained at least one plasmid sequence^22^. However, the microbial isolates are disproportionately represented by clinically relevant species, in which plasmid transmission is already known to be prevalent. In contrast, the vast majority of microbes remain uncultivated from globally relevant ecosystems, limiting our ability to sequence them and assess the ecological prevalence and functional contribution of MGEs, particularly in environmental contexts.

As an alternative, single-cell genome sequencing presents an opportunity to overcome this barrier by enabling high-resolution mapping of MGE host ranges directly within their native environmental context and at the cellular level^23,24^. The assumption underlying this approach is that the single cell that was sorted contained one or more MGEs within it or attached to cell surface materials (*e.g.* flagella). Here, we analyzed around 60,000 SAGs^23,25–32^, primarily derived from host-associated samples but including a range of environmental samples from soils, ocean, and freshwater. We assessed the prevalence of MGEs as a function of ecosystem type and evaluated MGE diversity and frequency in microbes from diverse lineages. We examined the host range of MGEs and investigated the genomic similarity both among MGEs sharing a host and among hosts sharing an MGE. Furthermore, by characterizing their transmission patterns and accessory genes, we explored MGE-mediated transmission of ARGs, and assessed the dissemination of defense-related and auxiliary genes in relation to MGE prevalence. Finally, we identified examples of unclassified circular elements with broad host ranges and high persistence across samples. Taken together, our results provide insights into MGE–host interactions, and highlight the potential of single-cell genomics to uncover genetic potential of mobile genetic elements across both host-associated and environmental microbiomes.

## Results

### MGE prevalence is ecosystem dependent

We assembled scaffolds from 55,839 SAGs collected from diverse ecosystems, comprising ocean water and sediment, human gut and oral cavity, and soil (Supplementary Table 1). We recovered 19,973 microbial genomes with >50% completeness and <5% contamination, and 117,596 plasmid scaffolds and 103,892 viral scaffolds that are greater than 2kb. MGE scaffolds were clustered 16,520 plasmid clusters and 25,340 viral clusters, respectively (see Supplementary Fig. 1 for an overview of the pipeline). The number of clusters may be overestimated, as many plasmid and viral genomes were fragmented and the fragments clustered separately. However, because each strain is often represented by multiple SAGs, overlapping and complementary coverage across SAGs helps reunite fragmented MGE contigs into the same clusters, thereby partially mitigating this bias. We verified that there was minimal cross-sample contamination by ensuring that near-identical scaffolds were not present in many SAGs from the same project (see Methods and Supplementary Fig. 2).

Prior to assessing the distribution of MGEs in microbial hosts, we quantified the prevalence of MGE clusters across different ecosystems, restricting the analysis to SAGs generated using identical single-cell processing and sequencing protocols (SAG-gel specifically), as different protocols (e.g. DNA amplification method) will significantly affect the MGE prevalence detected^33^. A few hundred unique SAGs were generated from non-host samples, and > 20,000 SAGs derived from host-associated ecosystems. We found that overall MGE prevalence is strongly influenced by both ecosystem type and MGE category (virus or plasmid). The majority of SAGs from host-associated ecosystems, specifically oral and gut, harbor viral and plasmid elements within individual cells (Fig. 1a and Supplementary Fig. 3). Viral clusters are found in ∼75% of human associated (gut and oral) bacteria, ∼45% of soil SAGs, ∼30% of ocean water and ∼30% of ocean sediment, whereas plasmid clusters occur in ∼70% of SAGs in all environments except ocean sediment (Fig 1). Viral and plasmid prevalence are broadly comparable across ecosystems, with the exception of ocean water, where plasmids occur at approximately three times the viral prevalence.

**Fig. 1.**
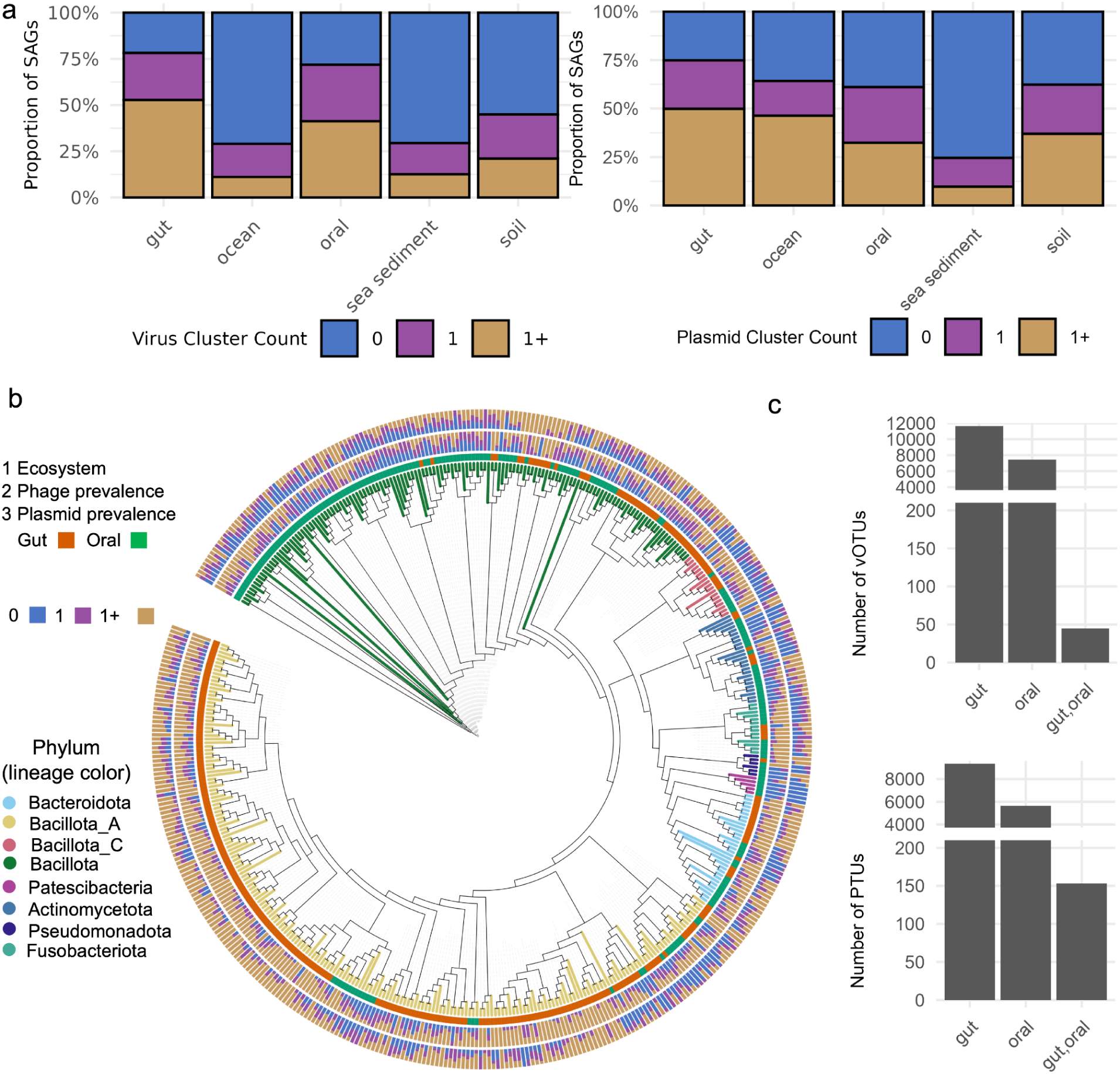
Prevalence of viral and plasmid clusters across diverse ecosystems based on single-cell genome sequencing. **a**, Proportion of single amplified genomes (SAGs) in each ecosystem containing 0, 1, or more viral or plasmid clusters per SAG. **b**, Same as (a), but restricted to SAGs derived from diverse species sampled from oral and gut ecosystems. **c**, Total number of viral and plasmid clusters identified in SAGs from the gut, oral, or both ecosystems.

### Comparing MGE incidence between oral and gut microbes

We analyzed >7,000 SAGs from ocean water^27^, found low virus prevalence in marine environments across species from 11 different phyla, although certain species of Pseudomonadota and Cyanobacteriota show slightly elevated prevalence (Supplementary Fig. 4). However, because a different protocol was used to generate these SAGs, their prevalence cannot be directly compared with that observed in oral and gut ecosystems.

As few non-host environment SAGs were generated using the same protocol as applied for host-associated SAGs (and probably only were recovered from dominant organisms), we could not compare virus and plasmid incidence patterns across different species. Thus, we compared SAGs from the gut and oral microbiomes, which are represented by over 5,000 and 10,000 SAGs respectively, and compared MGE prevalence across species that are represented by more than three SAGs in each ecosystem. Most of the selected species are exclusively found in either the gut or oral microbiome. High MGE prevalence is consistently observed across gut- and oral-derived SAGs from diverse species (Fig. 1b). Closely related (but distinct) species in the gut, such as those within Bacteroidota and Bacillota_A SAGs tend to harbor more MGEs than their oral counterparts. This trend also generally holds for the limited number of species represented in both gut and oral environments (Supplementary Fig. 5 and 6).

We quantified the number of viral operational taxonomic units (vOTUs) and plasmid taxonomic units (PTUs) present in the gut and oral microbiomes, as well as those shared between the two ecosystems (Fig. 1c). There were significantly more vOTUs than PTUs in both gut and oral environments. However, three times more plasmids than viruses were shared between gut and oral samples. This may indicate a broader host range of plasmids compared to viruses. Alternatively, plasmids may tend to associate more with organisms shared between gut and oral bacteria than viruses.

### Most MGEs have a narrow host range

MGEs-host ranges deduced from metagenomic studies are often broader than hosts ranges typically reported from laboratory experiments^16^. To address this possible discrepancy, we examined the host range of the vOTUs and PTUs using the MGE-host linkages established in this study. We calculated the minimum pairwise average nucleotide identity (ANI) for microbial host genomes harboring each MGE cluster (defined by sharing 95% ANI with ≥85% alignment fraction). In most cases, the microbial genomes hosting closely related MGEs shared an ANI above 97%. Using the well accepted ANI cutoff of 95% for bacterial species, these results suggest strain-level host specificity (Fig. 2a).

**Fig. 2.**
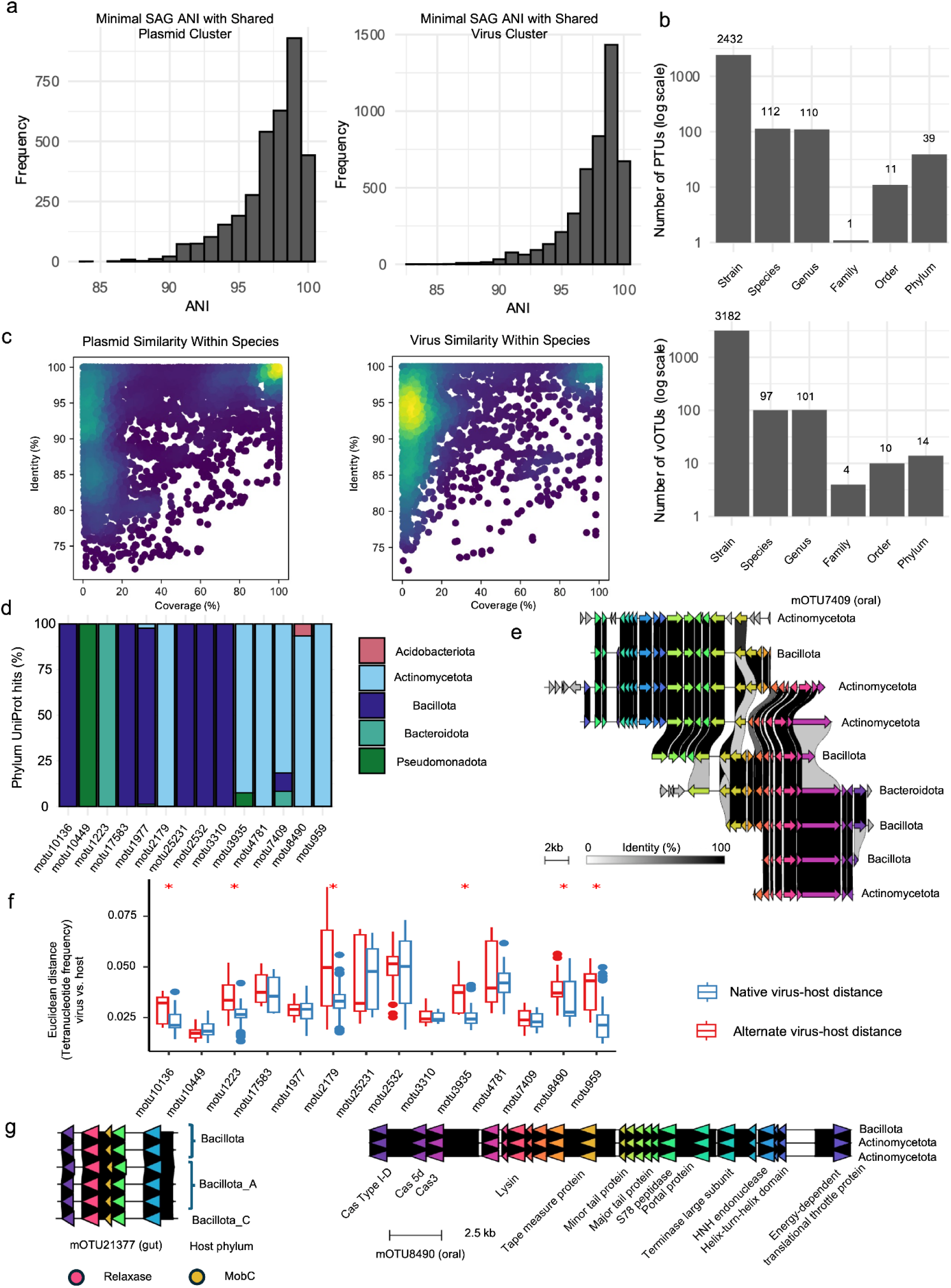
Host range of plasmid and viral clusters inferred from single-cell genome sequencing. **a**, Minimum average nucleotide identity (ANI) between SAGs that share the same plasmid or viral cluster. **b**, Host range of plasmid or viral clusters identified in more than two SAGs. **c**, ANI between plasmid or viral clusters found in SAGs of the same species. **d**, Phylum-level distribution of best UniProt taxonomic matches for all coding sequences in vOTUs exhibiting cross-phylum host range. **e**, sequence alignment of viral contigs from the vOTU mOTU7409. **f**, Comparison of primary and alternative virus–host tetranucleotide frequency distances across vOTUs with host range across phyla (See also Supplementary Fig. 8). For each ORF within every member in each vOTU, only the best UniProt hit was retained. The phylum-level taxonomy of these best-hit sequences was summarized as the percentage of ORF matches assigned to each phylum per vOTU. **g**, Examples of circular and complete plasmid (left) and provirus (right) sequences identified from SAGs belonging to different phyla.

Although most MGEs were essentially species specific, we observed a variety of vOTUs and PTUs with host ANI values <95%, suggesting a broader host range. Given that high-throughput ANI calculations are most unreliable for <82% ANI^34^, we also assessed the host range of MGE clusters containing more than two members (Fig. 2b) to understand broad range host range. As the host taxonomic rank increased, the number of shared MGE clusters generally decreased. However, we observed a gap at the order level, above which the number of diverse host vOTUs and PTUs increased again, including dozens of PTUs and vOTUs with host range across different phyla. We identified circular and complete plasmid and viral contigs in dozens of different SAGs belonging to different phyla (Fig. 2g and Supplementary Fig. 7), further suggesting the existence of viruses with broad host ranges.

A detailed examination of broad host range vOTUs and PTUs (those spanning different genera or higher taxonomic ranks) revealed that most possess a dominant host, with over 50% of their members associated with the same host lineage, and in some cases exceeding 90% (Supplementary Table 2). Among the vOTUs with host ranges spanning different species or higher taxonomic ranks (Fig. 2b), 51 (22.6%) vOTUs contained members annotated with plasmid hallmarks (Supplementary Table 3), including Type IV secretion systems, relaxases and plasmid segregation proteins. These comprised five vOTUs across species, forty across genera, two across orders and four across phyla. To further assess the cross-phylum host range observed in Fig. 2b, we examined the phylum-level UniProt matches for all open reading frames (ORF) in the 14 vOTUs (see Methods). Ten of the 14 vOTUs showed >99% of ORFs matching a single bacteria phylum (Fig. 2d), and five of the ten vOTUs have UniProt phylum hits that match the primary phylum assignment, defined as the phylum supported by more than half of the SAGs hosting the vOTUs. The remaining five vOTUs have UniProt phylum hits that match a single non-primary host phylum. Among the remaining four vOTUs, only mOTU7407 displayed a notable secondary signal, with ∼10% of UniProt hits consistently mapping to a second phylum. In particular, mOTU7407 yielded 81.7% of hits to Actinomycetota, 10% to Bacillota, and 8.2% to Bacteroidota, aligning with its three phylum-level host assignments (Fig. 2e). To assess differences in host association signals, we compared the Euclidean distances between tetranucleotide frequency (TNF) profiles of viral contigs and their host SAGs (see Methods), separating each vOTU’s primary host (native virus–host distance) from alternative hosts (alternate virus–host distance). For the cross-phylum broad host-range vOTU mOTU7407, the TNF distance did not differ significantly between primary and alternative host comparisons (Fig. 2f). However, when extending this analysis across all broad host-range vOTUs, we observed that alternative host distances were frequently and significantly higher than primary host distances (per-vOTU Wilcoxon tests; Supplementary Fig. 8).

### Genomic divergence separates MGE populations that infect the same host

We evaluated the genomic similarity of MGEs within clusters detected in SAGs from the same microbial species using ANI and aligned fraction (AF) metrics (Fig. 2c). The density distribution revealed two prominent hotspots for both vOTUs and PTUs: one at ANI >95% and AF >95%, corresponding to highly similar or nearly identical MGEs (i.e., of the same population); and another at ANI >80% but AF <20%, likely reflecting partial homologies.

We performed network analysis focusing on MGE clusters with inferred broad host ranges where each node represents a host species and each edge represents an MGE cluster shared between connected species (Fig. 3). In the plasmid network, we observed a distinct pattern of clustered MGE interactions within the phylum *Bacillota*, primarily driven by PTUs with host ranges restricted to the genus level (i.e., cross-species but within-genus). An exception was a hub species, *Streptococcus salivarius* (highlighted by a red ellipse), which harbored multiple PTUs connecting Bacillota with other phyla. Additionally, we identified several PTUs with broad host ranges spanning phyla, most notably linking Bacillota_A and Bacteroidota. Sporadic cross-phylum linkages were also observed between Actinomycetota and Bacillota. A similar pattern was observed for vOTUs, the majority of vOTUs within Bacillota exhibited narrow host ranges limited to within-genus associations. Again, *Streptococcus salivarius* emerged as a hub species, harboring vOTUs that bridge Bacillota with other phyla. Cross-phylum vOTUs were predominantly associated with hosts from Bacillota_A and Bacteroidota. The results suggest that broad host range MGEs are concentrated in certain taxa, with specific species such as *Streptococcus salivarius* acting as key hubs that connect distant hosts and potentially facilitate cross-phyla gene transfer.

**Fig. 3.**
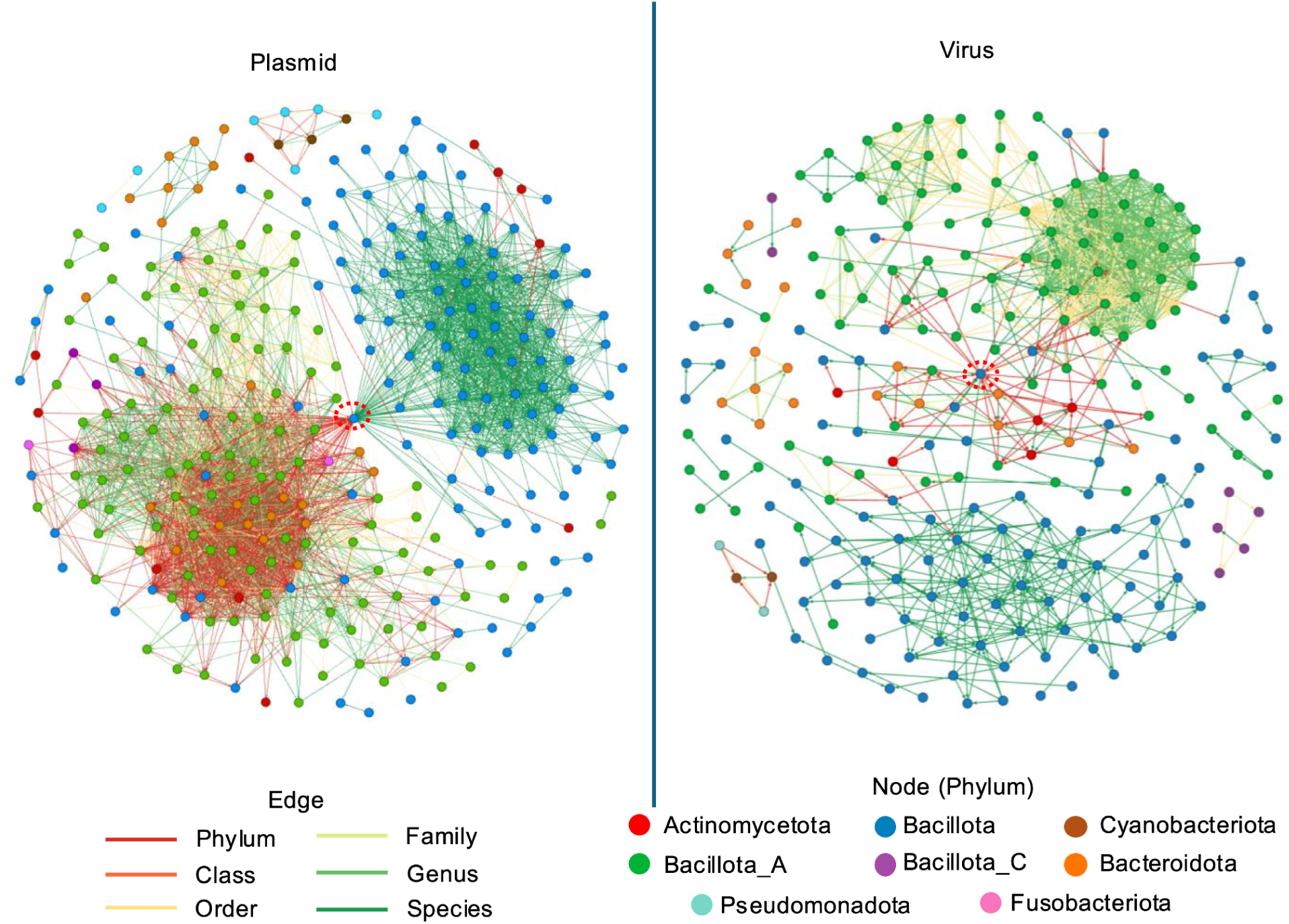
**Clustering of SAG hosts associated with broad-host-range plasmid and viral clusters.** Each node represents a microbial species, and an edge between two nodes indicates that the same mobile genetic element (MGE) is associated with both hosts. Edge colors denote the lowest common ancestor rank shared by the connected species. Node colors indicate the microbial phylum.

### MGE- and Host-Specific Horizontal Gene Transfer of ARGs

We identified plasmid- and virus-mediated HGT of ARGs, predominantly in host-associated ecosystems and rarely in other environments, with plasmids contributing more frequently than viruses (Fig. 4a), consistent with previous observations^35^. Based on these findings, we focused our analysis on MGE-mediated ARG transmission using oral and gut SAGs from Kawano-Sugaya et al.^23^. While the original study estimated that approximately 2% of ARGs were located in plasmids or phages, our investigation of ARG HGT revealed distinct transmission patterns and uncovered a much greater incidence of ARGs dissemination. The dominant mobile ARGs differed between the gut and oral environments, with blaR1 and erm(B) being the most prevalent in each, respectively. However, certain ARG classes, such as *mef(A)* and *msr(D)*, are carried by both phages and plasmids in comparable numbers. Potential ARGs transmissions are dominated by species in Bacillota and Bacillota_A (Fig. 4b). By distinguishing between recipients, a host cell carrying MGEs with ARGs, and donors, a host cell from which the ARGs in those MGEs likely originated, we observed that certain species acted as central hubs in ARG transmission, being prevalent in both donating and receiving roles. However, because these species were also among the most abundant in our dataset, their apparent centrality likely reflects a sampling artifact rather than true biological prominence. Therefore, these species were excluded from subsequent analyses. Among the less prevalent species, species like *Blautia_A faecis* (indicated by a blue arrow) emerge as notable hotspots for acquiring ARGs, having received ARGs from over 20 distinct donor species, while contributing to only a limited number of recipients. In contrast, *Thomasclavelia ramosa* (green arrow) stands out as a prominent ARG donor, transferring ARGs to more than 20 different recipient species, yet acquiring ARGs from only a few species.

**Fig. 4.**
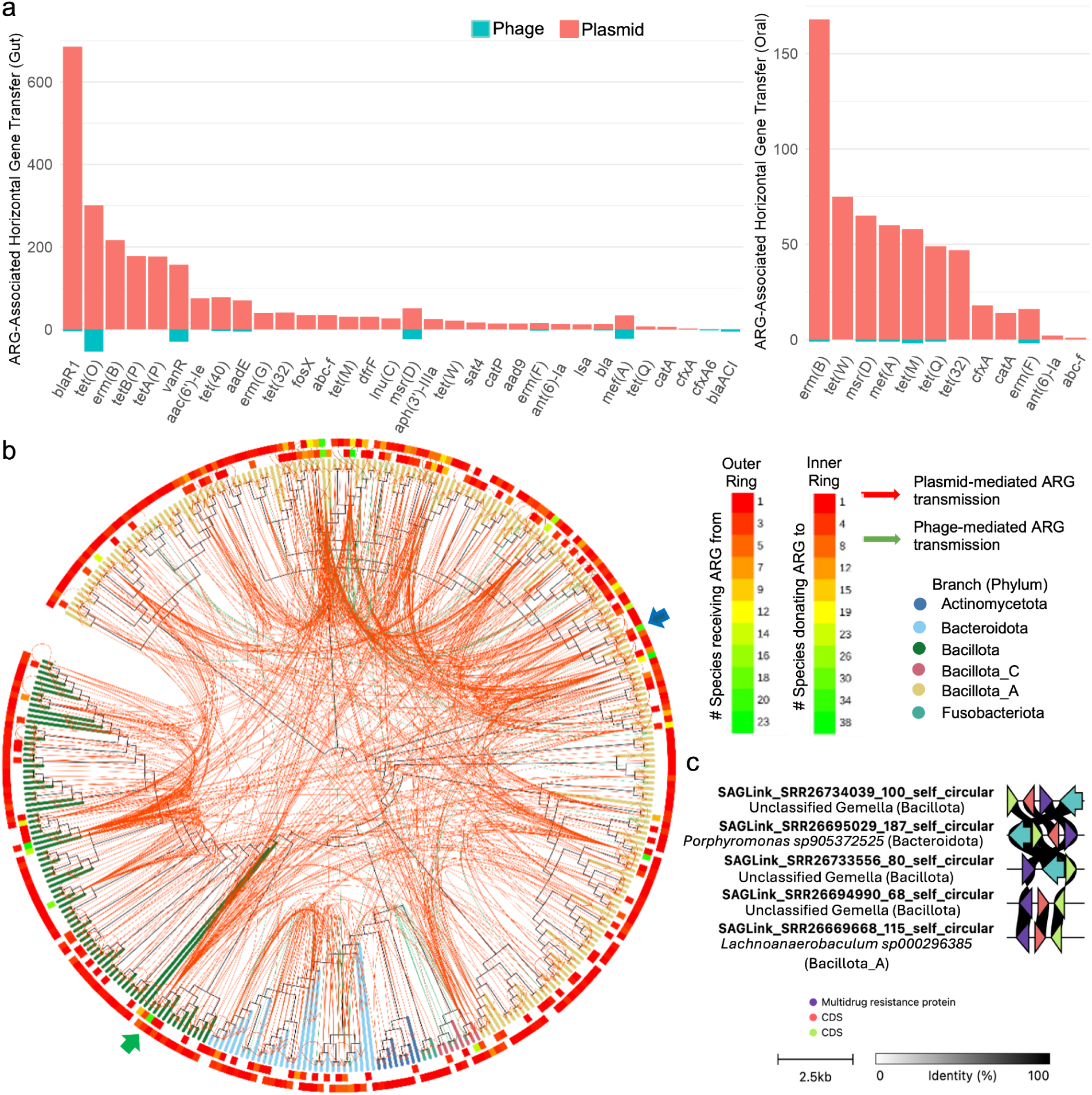
Plasmid- and virus-mediated transmission of antimicrobial resistance genes (ARGs) determined by single-cell genome sequencing. **a**, Prevalence of ARGs associated with plasmid- or virus-mediated transfer in gut and oral microbiomes. **b**, Inferred donors and recipients of plasmid- or virus-mediated ARG transmission events. **c**, An MGE with a broad host range spanning multiple phyla encodes a multidrug resistance protein.

### Unclassified Mobile Genetic Elements Exhibit Broad Host Range, Wide Distribution, and Harbor ARGs

MGEs have typically been identified using machine learning-based tools, which are inherently biased toward detecting known MGEs. However, extrachromosomal genetic elements such as Borgs^36^ and Mini-Borgs^37^, lack most canonical viral or plasmid hallmarks, yet appear to play important ecological and evolutionary roles. We identified an unclassified MGE with a broad host range spanning multiple bacterial phyla (Fig. 4c). The 2.3 kb circular element, manually validated to be complete (Supplementary Fig. 9), encodes a multidrug resistance protein and a transposase but lacks hallmark features of viruses or plasmids. This element was identified in SAGs from multiple phyla, including Bacillota, Bacteroidota, and Bacillota_A. We also identified a 6.4 kb circular element (Supplementary Fig. 10), manually verified to be complete (Supplementary Fig. 11). Although the element lacks canonical plasmid or viral hallmarks, the presence of a putative viral replication protein, a DNA segregation ATPase, and competence-associated proteins suggests it may utilize plasmid-like mechanisms for replication and maintenance, and is therefore likely to represent a cryptic plasmid. Notably, the element matched a CRISPR spacer identified in 12 single amplified genomes SAGs, all classified as *Anaerostipes hadrus* and originating from two unrelated projects, indicating that it is prevalent within this species.

### MGE-Enabled Dissemination of Defense and Metabolic Functions

In addition to investigating the horizontal transfer of ARGs, we also examined the MGE-mediated transfer of defense-related genes. To control for variation in MGE prevalence, we limited our analysis to SAGs generated using the SAG-gel method, consistent with our earlier analysis. We observed widespread transfer of defense-related genes via MGEs, with plasmids contributing more frequently than viruses. Notably, the prevalence of transferred defense genes appeared to correlate with typical microbial densities across ecosystems, being highest in the gut, followed by the oral cavity, and lowest in the ocean (Fig. 5a). We also assessed the distribution of adaptive (CRISPR-Cas) and innate (e.g., restriction-modification) immune systems across individual SAGs (Fig. 5b). Most SAGs from gut, oral, and ocean samples encoded at least one defense system, and a substantial fraction harbored CRISPR-Cas loci. However, ocean-derived SAGs notably lacked detectable CRISPR-Cas systems, consistent with earlier metagenomic surveys^38^. Of the 1,855 MGEs with CRISPR spacer matches, only 35 (1.8%) co-occurred within the same SAG harboring the corresponding spacers (Supplementary Table 4), suggesting that most spacer matches record past exposure events rather than ongoing infections.

**Fig. 5.**
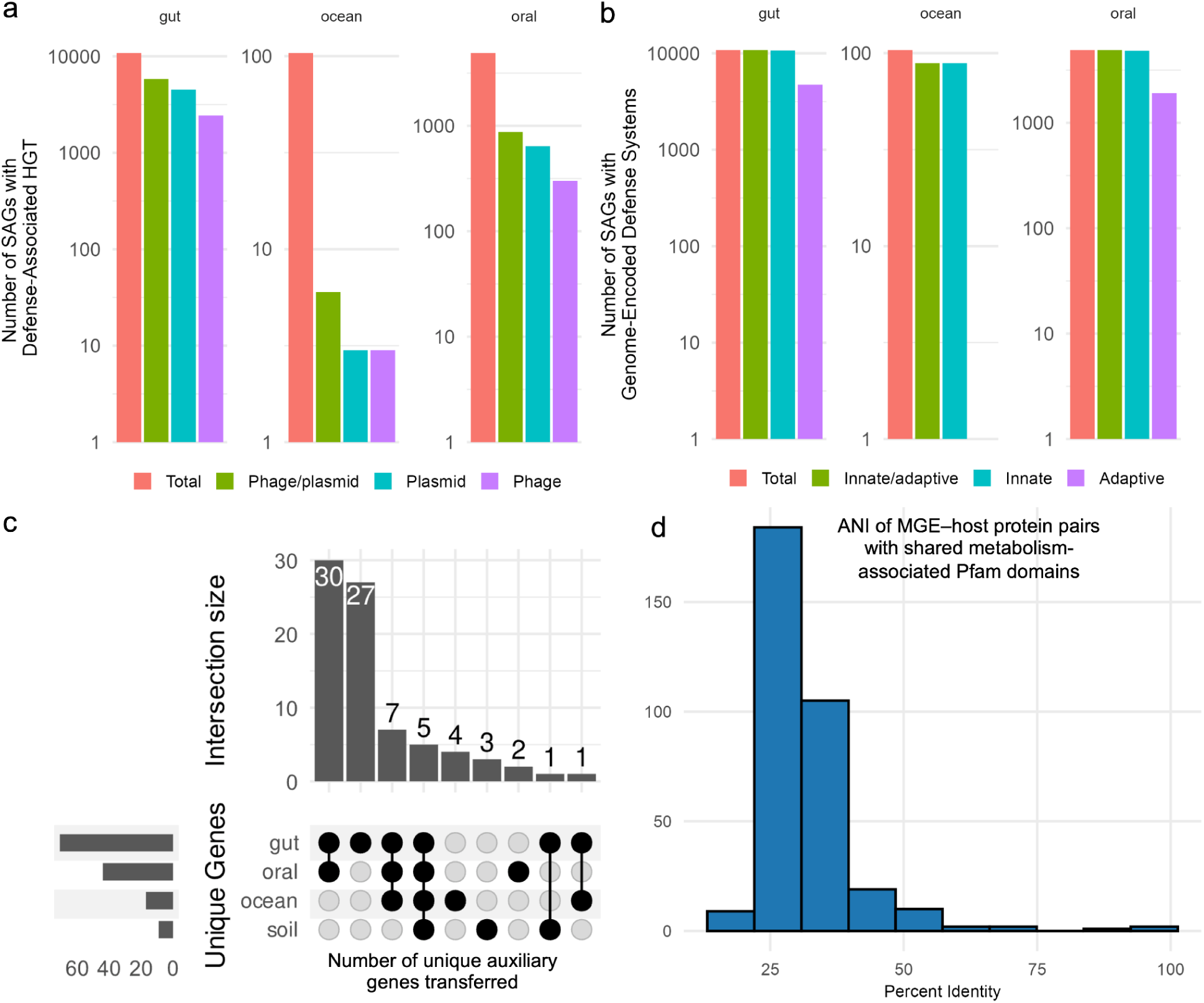
Defence systems encoded in SAGs or acquired via plasmid- or virus-mediated transfer. **a**, Number of SAGs containing defence-related genes horizontally transferred by plasmids or viruses in gut, ocean, or oral ecosystems. **b**, Number of SAGs with defence-related genes encoded natively in their genomes. **c**, Distribution of unique horizontally transferred auxiliary genes across ecosystems, showing genes shared among multiple environments or restricted to a single ecosystem. **d**, Amino acid sequence identity between pairs of mobile genetic element-encoded and host-encoded proteins that share the same Pfam domain.

**Fig. 6.**
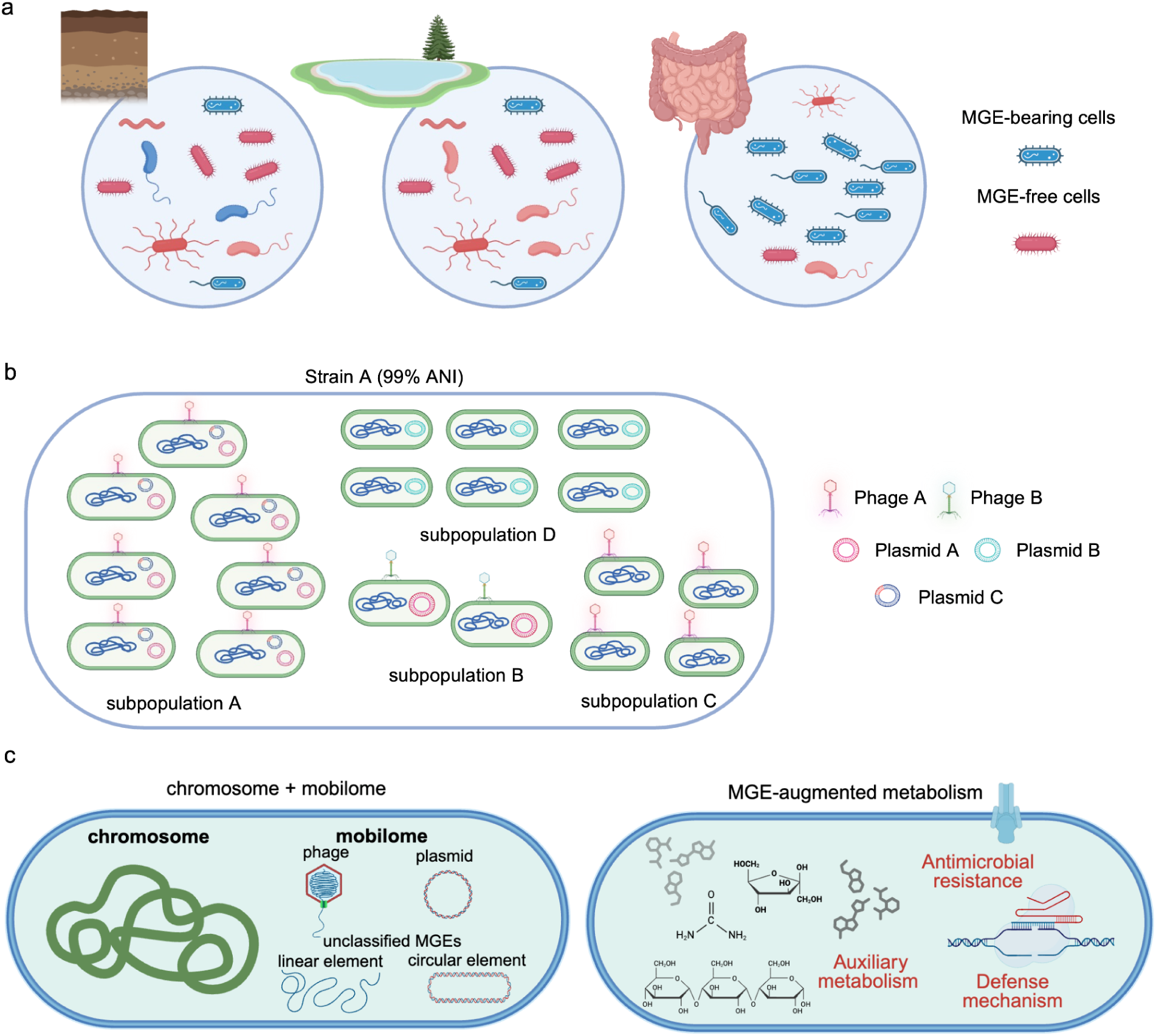
Conceptual model of MGEs shaping microbial functions and ecology across ecosystems. **a**, Microbial community composition in representative environments, highlighting the relative proportions of MGE-bearing and MGE-free cells. **b**, Despite high genomic similarity (ANI > 0.99), microbial strains can contain subpopulations with different MGEs. **c**, Schematic of the bacterial chromosome and the mobilome, including plasmids, phages, and other unclassified linear or circular MGEs. MGEs can augment host cell metabolism by providing genes related to antimicrobial resistance, auxiliary metabolic functions, or defense systems.

We identified a wide array of metabolism-associated genes transferred across all studied ecosystems (Supplementary Table 5). Notably, most of the horizontally transferred genes observed in the oral microbiome were also detected in the gut, suggesting frequent gene flow or a shared pool of mobile genetic elements between these two host-associated environments (Fig. 5c). In contrast, gene sharing between more phylogenetically or ecologically distant ecosystems, such as gut and soil, or gut and ocean was limited, with only a single HGT event identified in each case, consistent with expectations based on ecological separation and lower contact opportunities. Interestingly, several transferred genes appeared to be unique to specific environments, potentially reflecting ecosystem-specific selection pressures. For example, *thiosulfate sulfurtransferase* (K01011) and the *phosphate regulon sensor histidine kinase* (K07657) were uniquely identified in the ocean environment, highlighting the distinct nutrient cycling and regulatory challenges encountered in marine ecosystems (Supplementary Table 5).

We examined cases in which both the host and the associated MGE encoded the same function, identifying hundreds of instances where identical Pfam domains were present in both genomes (Fig. 5d). Examples of such shared domains included aminotransferases (PF00155.26), aldehyde dehydrogenases (PF00171.27), and phosphofructokinases (PF00365.25). Despite similarity at the domain level, amino acid sequence identity between MGE- and host-encoded proteins was generally low, with most pairs exhibiting less than 40% identity. This pattern is consistent with ancient acquisition events, acquisition from other hosts, or functional divergence over time. To broaden this analysis, we clustered all MGE- and host-encoded proteins. In cases where proteins encoded by the MGE and the chromosome of the same SAG clustered together, the majority (>99.9%) corresponded to genes expected to be in both (e.g., for replication, DNA modification, regulation).

## Discussion

Extensive metagenomic research has characterized the ecological and evolutionary roles of MGEs in shaping microbial communities, but the mechanisms have remained unresolved at the single cell level^9,39^. One of the fundamental unanswered questions is the prevalence of MGEs within individual microbial cells. While isolated genome sequencing has provided partial insights, it remains limited by loss of MGEs during growth in pure culture as well as cultivation biases. A recent analysis found that approximately 70% of gut microbial isolates, spanning over 72 genera, harbored at least one plasmid^22^. Notably, this prevalence varied by phylogeny: over 95% of *Faecalibacterium*, *Turicibacter*, and *Escherichia* isolates contained plasmids, significantly exceeding the average of 70% for isolates from gut microbiomes, but consistent with earlier findings in *E. coli*^21^. In contrast, the prevalence of both plasmid and prophage sequences is much lower in plant-associated microbial isolates, based on a survey of thousands of genomes from *Actinobacteriota*, *Bacteroidota*, *Firmicutes*, and *Proteobacteria*^40^.

Using single-cell genomics, we observed that approximately 75% of gut-derived SAGs carried at least one plasmid and one viral sequence, mirroring trends seen in isolated genome sequencing and corroborating the validity of our approach. Importantly, our analysis extends these findings to include uncultivated microbial lineages across seven phyla, revealing a general trend of elevated MGE prevalence in gut microbes. In environmental SAGs, MGE prevalence varied across ecosystems. Ocean-derived SAGs, for example, exhibited a higher frequency of plasmids compared to viruses. This may reflect both the low frequency of encounters between microbial cells and viruses in the ocean, and the reliance of viruses on host-to-host transmission for propagation. In contrast, plasmids can replicate autonomously within a single host and spread vertically through their hosts’ propagation. Moreover, plasmids often confer selective advantages, such as enhanced metabolic capabilities, which can be particularly beneficial in nutrient-limited environments like the ocean. It may also reflect the higher energetic cost of viral replication than plasmid carriage, as viral production requires capsid synthesis. The prevalence of MGEs appears to be shaped by both biotic and abiotic factors. This is illustrated by the case of SUP05 SAGs collected from varying ocean depths, where viral sequences were detected in roughly one-third of these SAGs^41^. The authors reported that viral sequence prevalence increased with depth, rising from 8% at 100 meters to 28% at 150 meters, and reaching 47% at 185 meters, coinciding with changes in environmental parameters such as oxygen and sulfur concentration^41^. Based on transmission electron microscopy, an average of 3.2% of heterotrophic bacteria and 1.5% of cyanobacteria in the ocean contain mature virus particles; considering that such assembled virions are only visible during the final ∼10% of the latent period, this translates to roughly 32% of heterotrophic bacteria and 15% of cyanobacteria being infected at any given time^42^. This level of infection is broadly consistent with our findings. To achieve a more comprehensive understanding, additional sampling from non-host-associated environments is needed. Although our results are largely consistent with findings from previous metagenomic surveys, microscopy observations, and isolate sequencing, they likely underestimate the true prevalence of MGEs in cells due to incomplete genome recovery during the whole-genome amplification process. Higher MGE prevalences have been reported in SAGs generated using newer alternative whole-genome amplification methods^33^; however, the number of such SAGs available to date remains limited.

Additionally, our results underscore that even among the same microbial species inhabiting different ecosystems, the prevalence of plasmids as well as viruses can vary substantially (Supplementary Figs. 5 and 6), highlighting the dynamic nature of MGE–host interactions (Fig. 6a). This pattern has also been observed in cultured isolates, where freshwater *E. coli* strains exhibited significantly lower plasmid frequencies compared to those isolated from the gut, accompanied by reduced frequencies of ARGs, virulence factors, and bacteriocins and higher frequency of capsule systems^21^. Additionally, we found that even within a single microbial strain, defined as sharing >99% ANI, distinct subpopulations could be identified based on their MGE content, highlighting intra-strain variation (Fig. 6b and Supplementary Table 6). For instance, in *Faecalimonas phoceensis*, one subpopulation of 87 SAGs carried a specific circular virus sequence, while another subpopulation of 90 SAGs carried a different one (Supplementary Table 6).

The prevalence of MGEs appears to influence the distribution of host defense systems. Ecosystems with lower MGE prevalence tended to contain fewer cells encoding adaptive immune systems, with gut-derived SAGs showing the highest frequency, followed by those from the oral cavity, and the lowest in ocean-derived SAGs (Fig. 5a). This pattern is consistent with recent metagenomic surveys, which similarly report that the abundance of phage defense systems varies across environments and generally increases with viral density^43^. The optimal growth temperature of microbes also influences the incidence of CRISPR–Cas systems, being lowest in psychrophiles and highest in hyperthermophiles^38^. Furthermore, nearly all gut and oral SAGs encoded at least one innate immunity system, whereas a substantially lower proportion of ocean SAGs carried innate immune defenses (Fig. 5b). The observed relationship between MGE prevalence and host defense systems may reflect a trade-off between the protective benefits of immunity and the fitness costs associated with replication, constitutive expression, and the risk of autoimmunity^44^. These dynamics are further shaped by factors such as MGE pressure, mutation rates, nutrient availability, and horizontal gene transfer. In ecosystems where MGEs are less prevalent, the metabolic burden of maintaining immune defenses may outweigh their selective advantages, reducing the evolutionary incentive to retain such systems. Interestingly, recent comprehensive analyses of CRISPR spacer sequences have revealed active, diverse, and rapidly turning over spacer repertoires^16^, suggesting that the absence of immunity in subpopulation may be mitigated by community-level immunity. However, broader and more systematic sampling of environmental SAGs is needed to better understand the dynamics of population scale immunity and its relationship to microbial phylogeny.

In addition to shaping population immunity, MGEs can augment host metabolism and adaptability by carrying and transferring genes related to antimicrobial resistance and auxiliary metabolic functions. We identified a diverse array of such auxiliary genes encoded by MGEs, many of which were horizontally transferred and enriched in host genomes (Fig. 5c, d). Given the widespread presence of MGEs across microbial cells from diverse ecosystems, we find that the metabolic capacity of an individual microbial cell should not be solely defined by its chromosomal content, but by the combined genomic repertoire of both the host genome and its mobilome (Fig. 6c).

We also found that the majority of MGE clusters exhibit a narrow host range, although broad-host-range MGEs remain common (Fig. 2). The number of MGEs with broad host ranges generally decreased with increasing taxonomic rank, except for a notable rise above the order level. The surge of vOTUs with a cross-phyla host range was also observed by Roux et al.^16^ by matching spacers with virus sequences. Isolated genome sequencing has shown that approximately 1.4% of plasmid clusters, defined at a 99% ANI threshold, from gut microbes spanning 72 genera can replicate in multiple host species^22^. A higher proportion of broad host range plasmids would be expected when using a lower clustering threshold (e.g., 95% ANI), as applied in our study. A large-scale analysis of over 10,000 reference plasmid sequences by Redondo-Salvo et al.^45^ found that 16% of plasmid clusters had host ranges spanning across bacterial families or higher taxonomic ranks, a frequency notably higher than observed in our dataset. This discrepancy may reflect the overrepresentation of gut-associated plasmids in reference collections, particularly those from clinical or medically relevant contexts. As for virus host range, Roux et al.^16^ leveraged CRISPR spacer matches from over 450,000 metagenomes to assign host taxa to viral sequences, reporting that 12.6% of viruses were predicted to infect hosts from multiple bacterial classes. This estimate, derived from unclustered viral genomes, may underestimate the true breadth at the vOTU level due to redundancy among closely related viral genomes.

Despite the predominance of narrow host range MGEs, we also identified a subset of MGEs with exceptionally broad host ranges spanning multiple phyla (Fig. 2, Fig. 3 and Supplementary Fig. 7). These vOTUs and PTUs with extraordinary host range may represent population-level host range expansion driven by factors such as receptor-binding protein variation, in which individual virions/plasmid typically still retain a narrow host specificity^46^. Nevertheless, our findings suggest that viruses and plasmids with identical genomic content may still infect a broad host range, potentially spanning across phyla (Fig. 2d and Supplementary Fig. 7). Similarly, several reports have documented cases where viruses infect hosts across different phyla, primarily based on methods such as CRISPR spacer matches^16,20,47,48^, but also by proximity-ligation-based (e.g. Hi-C) approaches^49^. Moreover, we found a 2.3 kb unclassified MGE encoding ARGs with broad host ranges, representing, to our knowledge, the first reported instance of MGEs spanning multiple phyla that carry ARGs. It is likely a circular replicative intermediate of a transposon that piggybacks on broad-host-range MGEs. Alternatively, it may represent a miniplasmid, as plasmids as small as 846 bp have been isolated and sequenced, lacking recognizable plasmid hallmark genes^50^. The absence of identifiable autonomous replication or conjugation genes, despite its presence across diverse taxonomic groups, suggests that it may be co-mobilized in association with other MGEs like conjugative helper plasmids.

Notably, inter-phyla host range was most frequently observed between Bacteroidota and Bacillota_A. Interestingly, *Streptococcus salivarius* emerged as a key hub species capable of mediating cross-phylum transfer for both plasmids and viruses, linking Bacteroidota, Bacillota, and Bacillota_A. In contrast, most other species within Bacillota harbored MGEs with predominantly narrow host ranges. *S. salivarius* is a widespread commensal bacterium found throughout the digestive tract, as well as on the skin, in breast milk, and other body fluids^51^. In addition, *S. salivarius* has been shown to be naturally competent^52^ and harbors a diverse repertoire of mobile genetic elements^53^. This highlights the potential role of *S. salivarius* as an important mediator of HGT across phyla in diverse human microbiomes, likely facilitated by its natural competence, ecological ubiquity, and association with diverse mobile genetic elements. Similarly, we identified hub species that acted either as major donors or as frequent recipients of ARGs in MGE-mediated HGT events across phylogenetically distant microbes (Fig. 4b). The results indicate that ARG transmission is not randomly distributed across taxa; rather, certain species emerge as key contributors to the horizontal spread of resistance, exerting a disproportionate influence on the resistome structure within their microbial communities.

Although viruses with exceptionally broad host range (e.g. across different phyla) are frequently reported through *in silico* analyses, to the best of our knowledge, only a single study to date has experimentally isolated viruses capable of infecting hosts across phyla^54^. Several factors may account for this discrepancy between computational predictions and empirical evidence. First, the apparent broad host range inferred *in silico* may reflect community-level rather than individual-level host range, as described previously. Second, most current MGE classifiers, such as geNomad, employ a binary framework that categorizes sequences strictly as plasmids or phages. This limitation can lead to the misclassification of hybrid elements, such as conjugative phage–plasmids that naturally exhibit broader host ranges as viruses (Supplementary Table 3). Consistent with our findings, marker gene–based analyses estimate that roughly 7% of sequenced plasmids and 5% of phages in the RefSeq database are in fact phage–plasmids^55^. Additionally, vOTUs and PTUs are generated by clustering viral and plasmid sequences separately. However, when we combined both viral and plasmid sequences and applied the same clustering pipeline, we observed instances where viruses and plasmids were grouped together, suggesting the presence of hybrid or mosaic elements that bridge these two categories. These findings highlight the need to analyse MGEs within a unified framework that captures the evolutionary continuum among mobile genetic elements including phages, plasmids, satellites, and other intermediates rather than treating them as discrete entities. A more modular classification scheme, as proposed by Land et al. ^56^, could better accommodate this diversity and should be considered in the development of future bioinformatic pipelines. Third, as noted by Roux et al. ^16^, CRISPR spacer matches may capture DNA uptake events (e.g. via conjugation, transformation and outer membrane vesicle) by non-host cells rather than productive infections. Likewise, physical association detected by single-cell genomics or Hi-C approaches may indicate viral entry or physical attachment without replication. We find that some broad host range vOTUs and PTUs showed a single dominant host comprising over 90% of their members (Supplementary Table 2 and Fig. 2d). This indicates that most associations are host-specific, with occasional but recurrent interactions across hosts potentially reflecting transient ecological events, such as DNA entry, rather than productive infections. Given these considerations, evidence for exceptionally broad host ranges is best interpreted through multiple, complementary lines of support, for instance, integrating physical associations with immune signatures such as CRISPR spacer targeting or shared methylation motifs between virus and host. Nonetheless, cell entry itself remains ecologically meaningful, as it exposes hosts to foreign genetic material, can activate immune defenses, and creates opportunities for gene exchange or latent persistence that influence microbial evolution. Moreover, the consistent presence of circular and complete plasmids and viruses across multiple, phylogenetically distinct SAGs (Fig. 2g; Supplementary Fig. 9) strongly suggests that these broad host range associations are biologically meaningful.

Finally, the narrow host range typically observed in cultured isolates may reflect skewed recovery of microbial lineages during laboratory cultivation and isolation. Replication and transfer capacities can be considerably broader under natural conditions. For example, IncP plasmids displayed markedly expanded host ranges when introduced into soil microbial communities, demonstrating that in the absence of physical or ecological barriers, broad-host-range plasmids can replicate across phylogenetically diverse bacteria^57,58^.

MGEs within the same species typically exhibit either partial homology or are near-identical sequences (Fig. 2c). The partial homologies potentially correspond to insertion sequences, transposons, intron-like elements, or other selfish genetic elements that contribute to MGE plasticity within species. This pattern expands upon similar observations previously reported for putative complete NCBI reference plasmids from members of the order Enterobacterales^45^. In plasmids, incompatibility groups describe sets of plasmids that cannot stably coexist in the same host cell because they share similar replication or partitioning systems, leading to competition. The low-AF and low-ANI relationships we observed for viral MGEs within the same species suggest that coexistence may similarly require viruses to be genomically distinct, possibly reflecting different replication strategies or machinery, and may also be influenced by phage-encoded superinfection exclusion mechanisms.

By integrating these observations into a unified conceptual framework (Fig. 6), we highlight the mobilome as a functional extension of the host genome, a conduit for cross-species and cross-ecosystem gene flow, and a key driver of microbial diversification and adaptation across ecological boundaries. Recently, Stepanauskas et al.^59^ found that HGT involves not only ecologically relevant “flexible” genes but also core genes by comparing the fraction of shared genes between SAG pairs against a lateral gene transfer–free model in marine prokaryoplankton. Importantly, ecologically significant transfer events were detected across order-level taxonomic distances, even though most HGT occurred among closely related cells. These findings largely align with our observations of MGEs exhibiting broad host ranges at deep taxonomic ranks, as well as our clustering results showing that MGEs and host proteins encompass both core and accessory genes. By directly analyzing the vectors of HGT, our results provide complementary evidence that MGEs are the principal drivers of gene flow across both shallow and deep evolutionary scales. Whereas Stepanauskas et al. ^59^ inferred transfer from gene sharing patterns, our single-cell MGE-centric framework identifies the specific vehicles and host contexts underlying these events. This highlights how a small subset of broad host range MGEs and hub species disproportionately shapes microbial community structure and evolution.

One of the major limitations of this study is the underrepresentation of SAGs from diverse ecosystems, largely due to the labor-intensive process of sampling, isolating, and sequencing of individual cells^60^. Microfluidic-based single-cell sequencing, which enables the simultaneous sequencing of tens of thousands of SAGs in a single run, has opened new opportunities to scale the exploration of MGE–host associations^32,61^. However, the widespread adoption of this approach has been limited by the need for custom-built instrumentation and specialized technical expertise. Recently, alternative microfluidic workflows that adapt commercially available devices and require minimal, standardized laboratory procedures have achieved comparable scalability for SAG generation, representing a more accessible and broadly applicable platform technology^62^. Alternatively, recent technological innovations have enabled targeted exploration of specific MGEs and their hosts through the use of multiplex barcoding PCR^63^.

In conclusion, our findings underscore the intricate and context-dependent nature of MGE–host interactions, which are shaped by ecological gradients, host phylogeny, and microbial defense systems. The identification of unclassified MGEs with broad host ranges, wide environmental distributions, and carriage of ARG expands the known diversity of the mobileome and emphasizes the need for detection approaches that extend beyond hallmark-based criteria. Collectively, these results demonstrate that ecological, phylogenetic, and defensive factors jointly govern MGE–host dynamics, reveal hallmark-elusive MGEs that broaden the mobileome landscape, and support a perspective in which cells and their MGEs function as an integrated genetic unit in nature.

## Methods

### Public single cell genomes sequencing processing

When available, raw reads from publicly available single-cell genome sequencing projects were downloaded from NCBI using the SRA Toolkit (https://github.com/ncbi/sra-tools). Assembled SAGs without associated raw reads were obtained from GORG^27^. A complete list of datasets and associated metadata is provided in Supplementary Table 1. Raw reads were trimmed for adapters and quality using BBDuk^64^ and sickle^65^, then merged using BBMerge^66^. Both merged and unmerged paired-end reads were assembled with SPAdes^67^ using the flags --sc --careful. Scaffolds longer than 500 bp were extended and assessed for circularity using COBRA^68^. The resulting COBRA-processed scaffolds from each SAG were used for downstream analyses. To assess potential sources of contamination in the single-cell sequencing projects, we examined the prevalence of individual scaffolds across SAGs within each project using skani triangle^34^. Specifically, for a given scaffold, we counted the number of SAGs containing scaffolds with an aligned fraction greater than 20% and ANI exceeding 99% relative to the query scaffold. In a single-cell sequencing project, the presence of contamination, such as those from reagents or environmental sources would result in DNA from the contaminant being detected across multiple SAGs. In all sequencing projects except PRJNA776656, the majority of scaffolds were present in less than 1% of SAGs, suggesting minimal cross-SAGs contamination. In contrast, SAGs from PRJNA776656, which includes SAGs from only two species contained a substantial proportion of scaffolds with >50% prevalence, validating the effectiveness of the detection method (Supplementary Fig. 2).

### MGE identification and clustering

Viruses and plasmids were identified from scaffolds using geNomad^69^. To detect potential unclassified MGEs, two additional strategies were employed: (1) scaffolds containing direct or inverted terminal repeats (DTR/ITR) were identified using the complete_genomes.py script from CheckV^70^ and filtered out if the average k21 frequency of the repeat exceeded 1.5, the repeat appeared more than twice on the scaffold, or the most frequent base in the repeat had a frequency >0.75; and (2) circular scaffolds not classified as viruses or plasmids by geNomad were retained as additional MGE candidates. All virus, plasmid, and potential non-canonical MGE sequences were further analyzed using VirSorter2^71^ and Vibrant^72^ and CheckV. As the majority of viral sequences identified by geNomad were also detected by VirSorter2 and VIBRANT, geNomad-derived MGEs were used for downstream ecological analyses, and these sequences were removed from the corresponding SAGs to retain MGE-free bacterial genomes. Proviruses were identified from excised virus contigs as well as viruses with provirus hallmarks (such as integrase and repressor protein) as identified by VIBRANT. Identified MGEs larger than 3 kb, were clustered into vOTUs and plasmid PTUs, respectively, using anicalc.py and aniclust.py from the CheckV, based on ANI >95% and aligned fraction >85%. The clustered MGEs were used for ecological analyses.

### SAG microbial genome analysis

After removing MGEs identified by geNomad from each SAG, the quality and taxonomy of the remaining microbial genomes were assessed. CheckM2^73^ was used to estimate completeness and contamination, and only SAGs with contamination <5% and completeness at least 10 times greater than the contamination were retained. For SAGs with completeness >50%, taxonomy was assigned using GTDB-Tk v2 with the GTDB R214 reference genome database^74^. For lower-completeness SAGs, taxonomy was determined using skani (skani search) against a pre-sketched GTDB R214 reference database (https://github.com/bluenote-1577/skani/wiki/Pre%E2%80%90sketched-databases). Only taxonomic assignments with an aligned fraction >10% and ANI >97% were retained (see following benchmark).

### Benchmarking skani for SAGs taxonomy classification

Since most of the SAGs have completeness below 50%, GTDB-Tk v2, which relies on the placement of individual marker genes, may not provide reliable classifications for these low-completeness genomes. Given the observed 95% ANI gap and the pattern of high genome coverage within species versus low coverage between species, we hypothesized that ANI and genome coverage against GTDB references could inform SAG taxonomy assignment. To test the hypothesis, we first used skani against the 5497 SAGs generated from a mock community of 4 known species from Zheng et al.^32^. Using skani as described above, we found that for SAGs with low reference coverage (<5%), even ANI values as high as 99% were insufficient for accurate species identification (Supplementary Fig. 12a). To test the minimal aligned fraction threshold required for accurate species classification, we simulated reads from 529 diverse reference genomes newly added in GTDB R220 (Supplementary Table 7) using InSilicoSeq^75^ at varying coverage and depth. After quality control and assembly as described previously, we generated 3,537 SAGs with a range of genome coverage levels. We again used skani to assign taxonomy and found that an aligned fraction >10% and ANI >97% were sufficient for accurate species identification (Supplementary Fig. 12b).

### MGEs and SAGs annotations

Open reading frames (ORFs) in MGEs and MGE-free SAGs were identified using pyrodigal-gv and pyrodigal, respectively, from the Pyrodigal package^76^. The ORFs were then annotated using a series of tools and databases, including Pfam^77^, KofamScan^78^, TIGRFAM^79^, and Rfam^80^. The defense and anti-defense mechanisms were annotated with PADLOC^81^ and DefenseFinder^82^. The spacers were identified from SAGs using MinCED^83^ after masking low-complexity regions. The identified spacers greater than 24 bp were searched against identified MGEs and non-cannonical MGEs using BLAST^84^ and only the hits with less 1 mismatch, 0 gap, and 100% spacer coverage were retained. ARGs were annotated using AMRFinderPlus^85^. tRNAs were annotated using tRNAscan-SE 2.0^86^. Using the genome annotations,

### Non-canonical MGE recovery and genome validation

We identified nearly identical circular non-canonical MGE sequences (with 100% coverage and >99% ANI) present in SAGs belonging to different phyla. We manually verified the circularity and genome integrity of these sequences using Geneious Prime 2022.2.2 (https://www.geneious.com). Additionally, we analyzed and visualized gene content and synteny among these MGEs using Clinker^87^ and Proksee^88^, based on GenBank files generated with Prokka^89^. We also predicted the structure of the protein from the non-canonical MGEs using AlphaFold2 via LocalColabFold with default parameters^90,91^. Structural homologues of the predicted structures were identified using Foldseek against AlphaFold Swiss-Prot database^92^.

### Host association and cross-ecosystem dynamics of MGEs identified from SAGs

By linking clustered MGEs to the SAGs from which they originated, we were able to assign MGEs to their hosts at the SAG level. When species-level classifications of the SAGs were available, they enabled MGE host associations to be resolved at the species level. To assess MGE prevalence across different ecosystems, we focused on comparing results from studies that employed identical single-cell processing. Specifically SAG-Gel from Chijiiwa et al.^29^, Nishikawa et al.^26^ and Kawano-Sugaya et al.^23^. This controlled comparison was necessary because the single-cell workflow, particularly the choice of whole-genome amplification protocol, has a substantial impact on MGE recovery, as demonstrated by benchmarking in Bowers et al.^33^. To compare MGE prevalence in the ocean using the SAGs reported by Pachiadaki et al. ^27^, we included only those SAGs sorted with SYTO-9, while excluding SAGs sorted with RSG, which specifically targeted live cells. To visualize MGE prevalence among species across different ecosystems, we selected species represented by more than three SAGs within each ecosystem. Most species were unique to a single ecosystem. To represent their phylogenetic relationships, we constructed a tree based on concatenated single-copy marker genes from GTDB reference genomes, using the gtdbtk infer function from GTDB-Tk v2. MGE prevalence for each species was then overlaid onto the phylogenetic tree and visualized using iTOL^93^.

We calculated pairwise genomic similarities, ANI and aligned fraction between MGEs and SAGs using skani within each project, applying the --small-genomes flag for MGEs but not for SAGs. Based on these metrics, we quantified the genomic similarity among SAGs harboring each MGE cluster and for each MGE cluster and minimal ANI among the SAGs. Similarly, within each species, we measured the genomic similarity among MGEs assigned to that species. We quantified the number of MGEs exhibiting host ranges across different taxonomic levels or at different ANI thresholds. For vOTUs and PTUs exhibiting host ranges above the strain level, only taxonomic assignments represented by at least two members were retained. For the 14 vOTUs exhibiting host ranges across different phyla, we examined the UniProt taxonomic profiles of all coding sequences in each vOTU member. Phylum level matches were summed per vOTUs following the approach described in Al-Shayeb et al.^94^. To visualize the host range of broad-host MGEs, those associated with hosts from different species or higher taxonomic ranks, we constructed a network in which each node represents a host species, and each edge represents an MGE shared between species, connecting the corresponding nodes. The network was visualized using Gephi (version 0.10.1)^95^. Since the majority of SAGs and diverse-host MGE clusters originated from gut and oral samples, we restricted the network analysis to MGE-host linkages established within these two ecosystems.

### Tetranucleotide frequency (TNF) analysis and virus–host distance measurements for the broad host range vOTUs

TNF profiles were computed for all viral contigs and host MGE-free host SAGs. Virus–host similarity was quantified as the Euclidean distance between TNF profiles. For each viral contig, the primary host was defined relative to that contig as the SAG assigned to its own host taxonomic group. SAGs belonging to other taxonomic groups, serving as primary hosts for other viral contigs within the same vOTU were considered alternative hosts for that contig. To assess whether viral contigs were compositionally more similar to their primary hosts than to alternative hosts taxonomic group, we performed per-vOTU one-sided Wilcoxon rank-sum tests. For each vOTU and each host-range rank, Euclidean TNF distances to primary hosts were compared with distances to alternative hosts. The test was conducted using a one-sided alternative hypothesis that TNF distances to alternative hosts are greater than TNF distances to primary hosts. Only vOTUs with at least one primary and one alternative comparison were included. When appropriate, *p* values were adjusted for multiple testing using the Benjamini–Hochberg procedure, and statistical significance was defined as *p* < 0.05.

### Detection and ecosystem-level comparison of MGE-driven ARG and defense gene transfer

We first identified MGE proteins that shared >99% amino acid identity and >99% alignment coverage (both query and target) with proteins encoded in MGE-free SAGs. To detect potential HGT events, we then examined the 5 kb flanking regions on both the 5′ and 3′ ends of the corresponding SAG sequences for the presence of HGT-associated features, such as tRNA genes, transposases, integrases, and phage or plasmid hallmarks, given that the mobility of ARGs is often mediated by transposases, integrases, and integrons^96^. Only cases where at least one HGT signature was present in the flanking region were retained as candidate HGT events. This criterion ensures that the events we report are high-confidence and mechanistically interpretable, minimizing the risk of misclassifying vertically inherited or assembly-derived sequences as HGT. While this approach likely underestimates the true extent of ARG mobility, particularly homologous recombination events where MGE signatures have been lost, it avoids the risk of inflating HGT counts through spurious identity-based matches. For example, a gene that is actually vertically inherited but highly conserved may be misclassified as an HGT event if it happens to be located near a residual MGE fragment. The majority of ARG-related HGT events were identified in host-associated datasets, specifically from SAGs generated in Kawano-Sugaya et al.^23^. Since SAG abundance reflects the underlying community structure, we leveraged this information to quantify ARG transmission across different MGEs and host species. To visualize these transmission patterns, we constructed a phylogenetic tree of species involved in ARG-associated HGT, as previously described. Using iTOL, we represented each HGT event as an arrow connecting the donor and recipient species, with the arrow pointing toward the ARG recipient.

We also compared MGE-mediated HGT of defense-related genes, based on annotations from PADLOC and DefenseFinder across different ecosystems. For this analysis, we focused on SAGs generated using the SAG-gel, as variations in single-cell processing methods can influence MGE recovery and, consequently, the observed prevalence of MGE-mediated HGT. In addition, we compared the overall defense system content encoded in SAGs, among those generated with SAG-gel.

### Identification of shared protein clusters between MGEs and hosts

All predicted proteins from MGEs and their corresponding host SAGs were combined into a single protein dataset and clustered using MMseqs2 easy-cluster ^97^. Clustering was performed with a minimum sequence identity threshold of 35% across 80% of the aligned region (--min-seq-id 0.35 --cov-mode 1 -c 0.8). Clusters containing both host- and MGE-encoded proteins from the same SAG were extracted for functional annotations using the methods described above.

### MGE Prevalence Within Microbial Strains

To assess MGE prevalence within microbial strains, we focused on strains defined by a shared MGE-free genome with >99% ANI. We examined the presence of MGEs across individual cells (SAGs), restricting our analysis to circular plasmids and phages. Only SAGs with >50% completeness and species represented by at least 50 SAGs within a dataset were included. MGE presence in each SAG was determined using CoverM^98^ with the parameters --min-covered-fraction 0.5 and --min-read-percent-identity 95.

## Data and code availability

The collected single-cell genome sequencing reads are publicly available and are summarized in Supplementary Table 1. The assembled sequences and identified MGEs reported in this study are available at Zenodo (DOI: 10.5281/zenodo.17994994).

## Acknowledgements

We thank the generators of published single cell amplified genome data that was instrumental to this study. This work was supported in part by Lyda Hill Philanthropies, Acton Family Giving, the Valhalla Foundation, Hastings/Quillin Fund - an advised fund of the Silicon Valley Community Foundation, the CH Foundation, Laura and Gary Lauder and Family, the Sea Grape Foundation, the Emerson Collective, Mike Schroepfer and Erin Hoffman Family Fund - an advised fund of Silicon Valley Community Foundation, the Anne Wojcicki Foundation through The Audacious Project at the Innovative Genomics Institute. MY, JFB, and RS were supported by Innovative Genomics Institute. We thank Leylen Miloslavich and Tasha Kayatsky for their laboratory support, and Drs. Lingdong Shi and Shuai Wang for insightful discussions. We also thank Glen Otero for computational infrastructure support.

## Authors contributions

RS conceived the study. MY, RS, and JFB designed the study. Bioinformatics analysis was performed by MY with guidance from RS. The manuscript was written by MY with input from JFB and RS. JFB and RS provided funding support.

## SUPPLEMENTARY FIGURES

**Supplementary Fig. 1.**
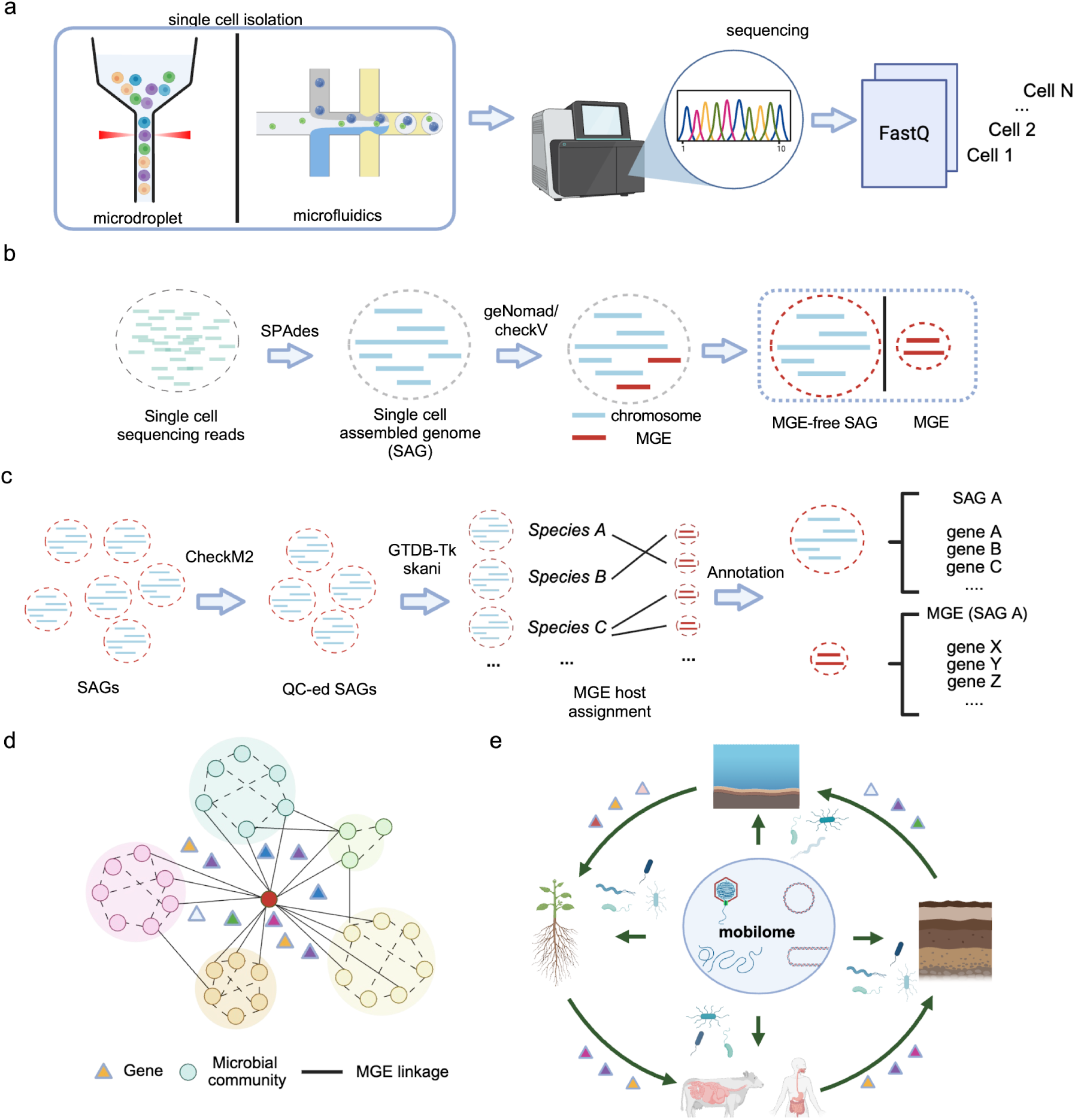
Overview of the single-cell genomics pipeline used. **a**, Data collected from single-cell isolation using microdroplet or microfluidics platforms, followed by whole-genome amplification, sequencing, and generation of per-cell FASTQ files. **b**, Scaffolds are assembled from single-cell data, MGEs are identified and separated from host chromosomes, producing both MGE-free host genomes and corresponding MGEs. **c**, Host genomes are taxonomically classified, MGEs are linked to their microbial hosts, and gene content is annotated for both host genomes and MGEs. **d**, Overview of downstream network analyses to trace MGE-mediated gene sharing within and between microbial communities, highlighting potential inter- and intra-population transfer patterns. Genes (triangles) are shared across different microbial communities (colored circles), with solid and dashed lines indicating potential inter- and intra-population gene transfer. **e**, Summary representation of ecosystem-scale analyses linking the mobilome to horizontal gene transfer across ecosystems, illustrating inferred routes of MGE exchange. Arrows indicate potential routes of MGE-mediated gene exchange.

**Supplementary Fig. 2.**
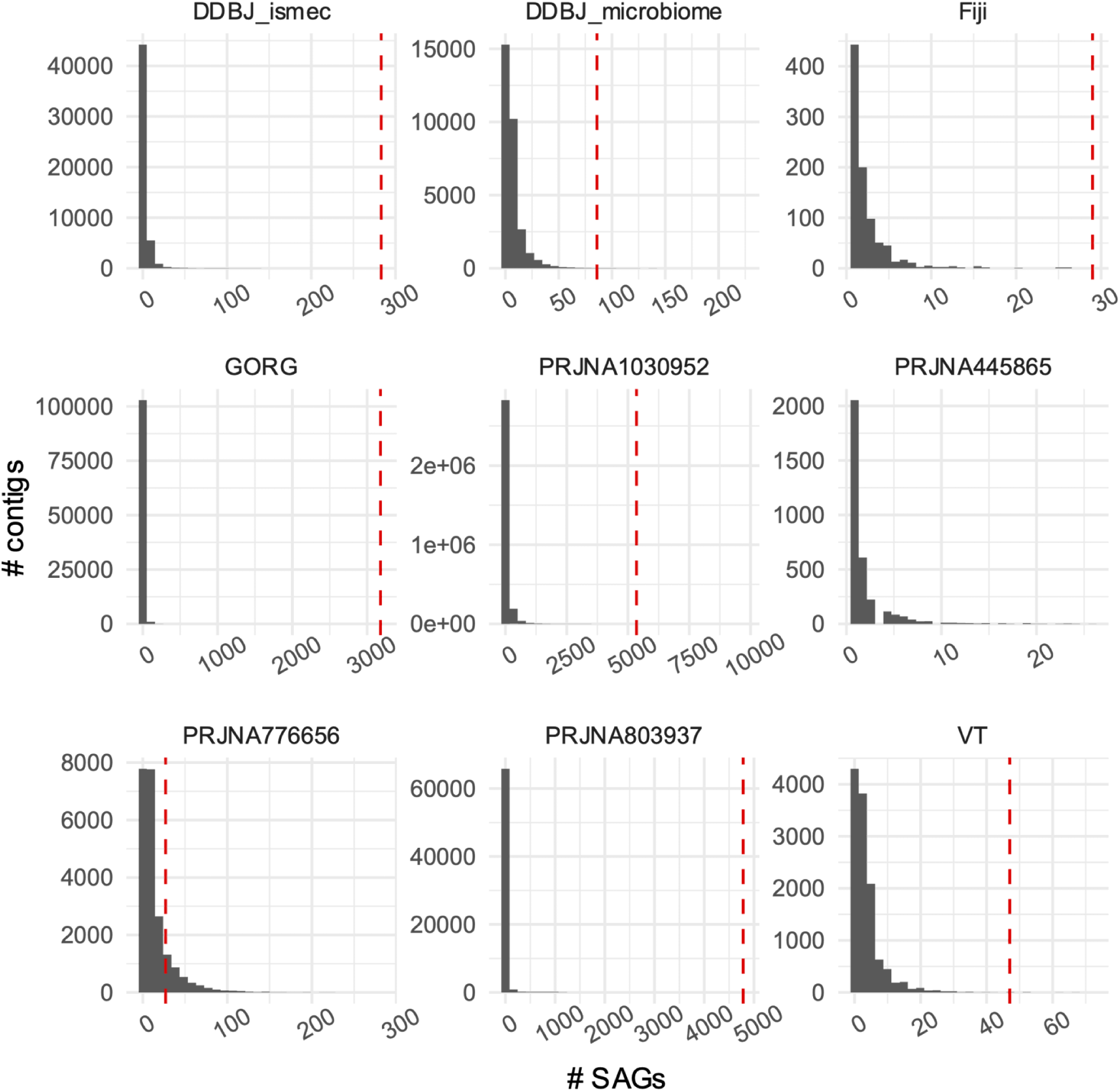
Prevalence of contigs across SAGs within the included sequencing projects.

**Supplementary Fig. 3.**
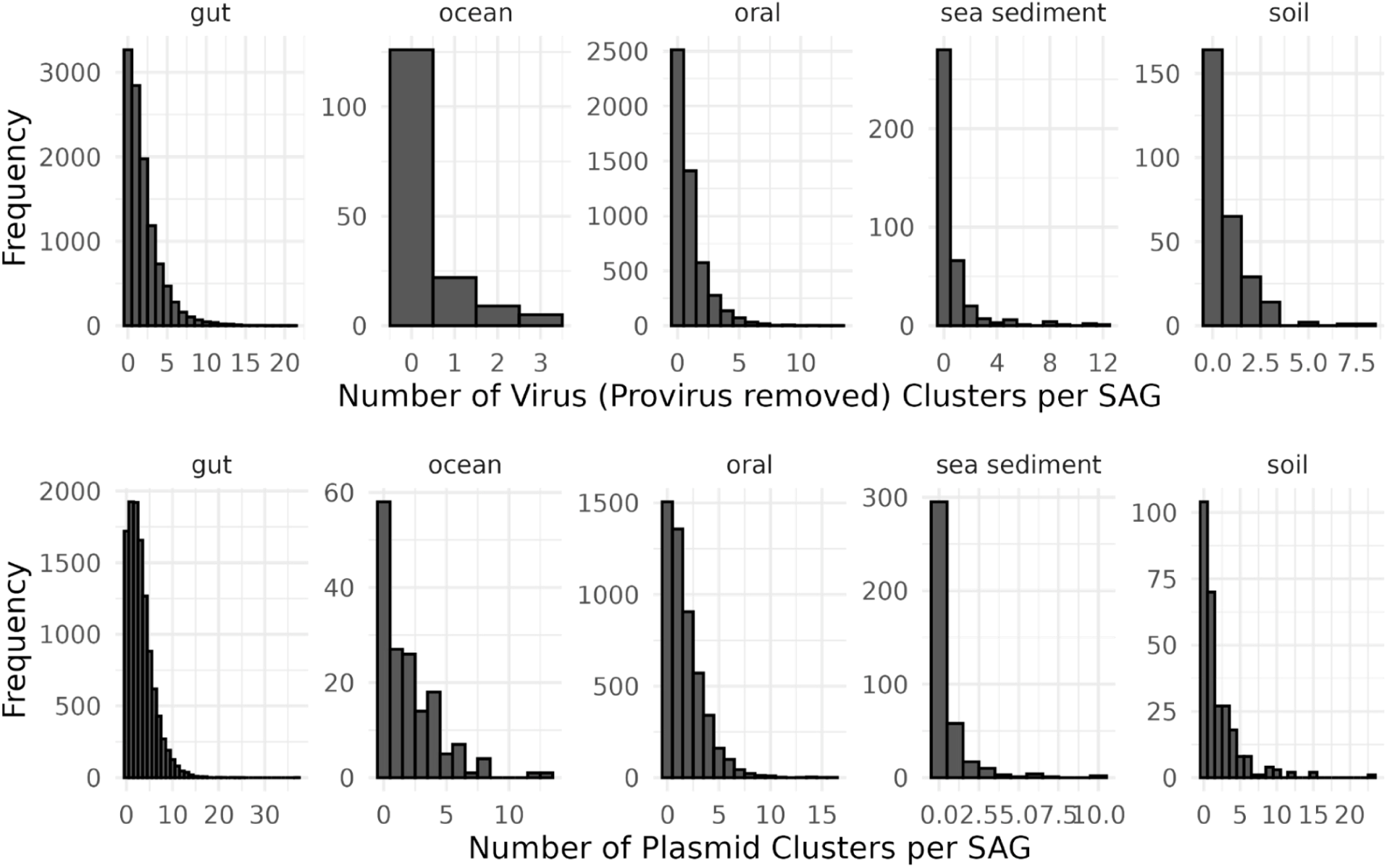
Number of clustered plasmids and viruses per SAG across various ecosystems.

**Supplementary Fig. 4.**
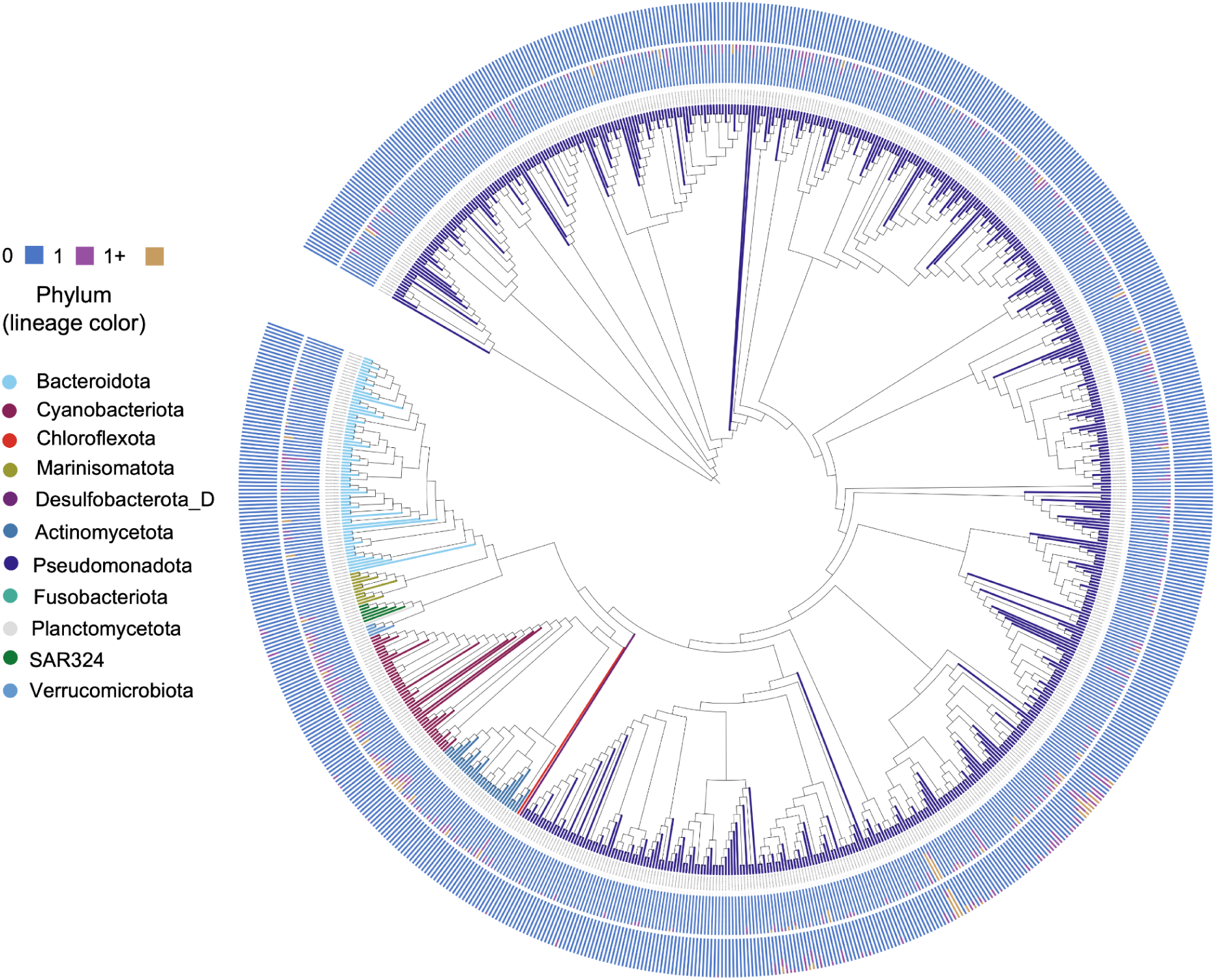
Prevalence of viral and plasmid clusters found in the ocean based on single-cell genome sequencing. Proportion of SAGs in each ecosystem containing 0, 1, or more viral or plasmid clusters per SAG.

**Supplementary Fig. 5.**
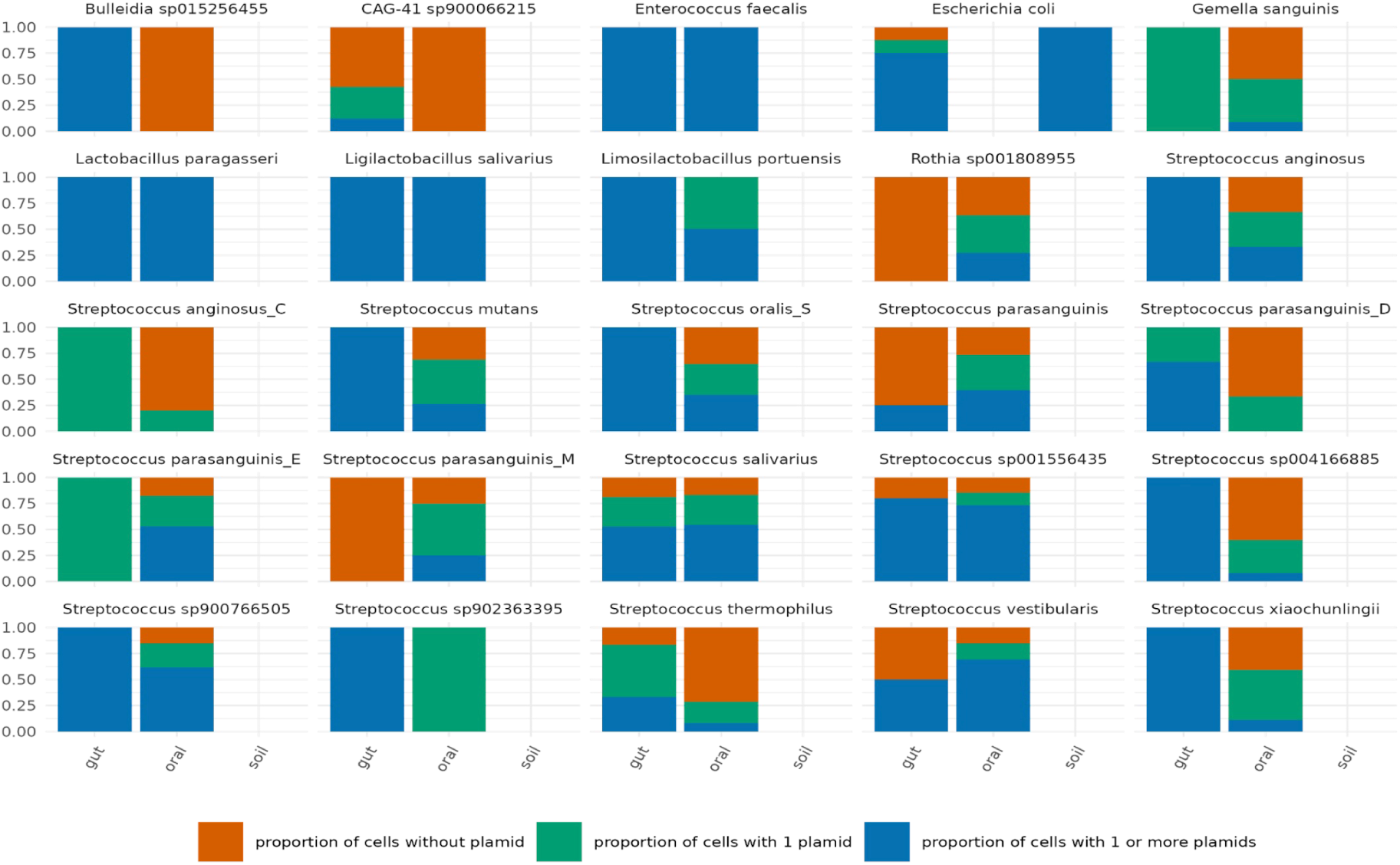
Prevalence of viral clusters shared among identical host species across diverse ecosystems.

**Supplementary Fig. 6.**
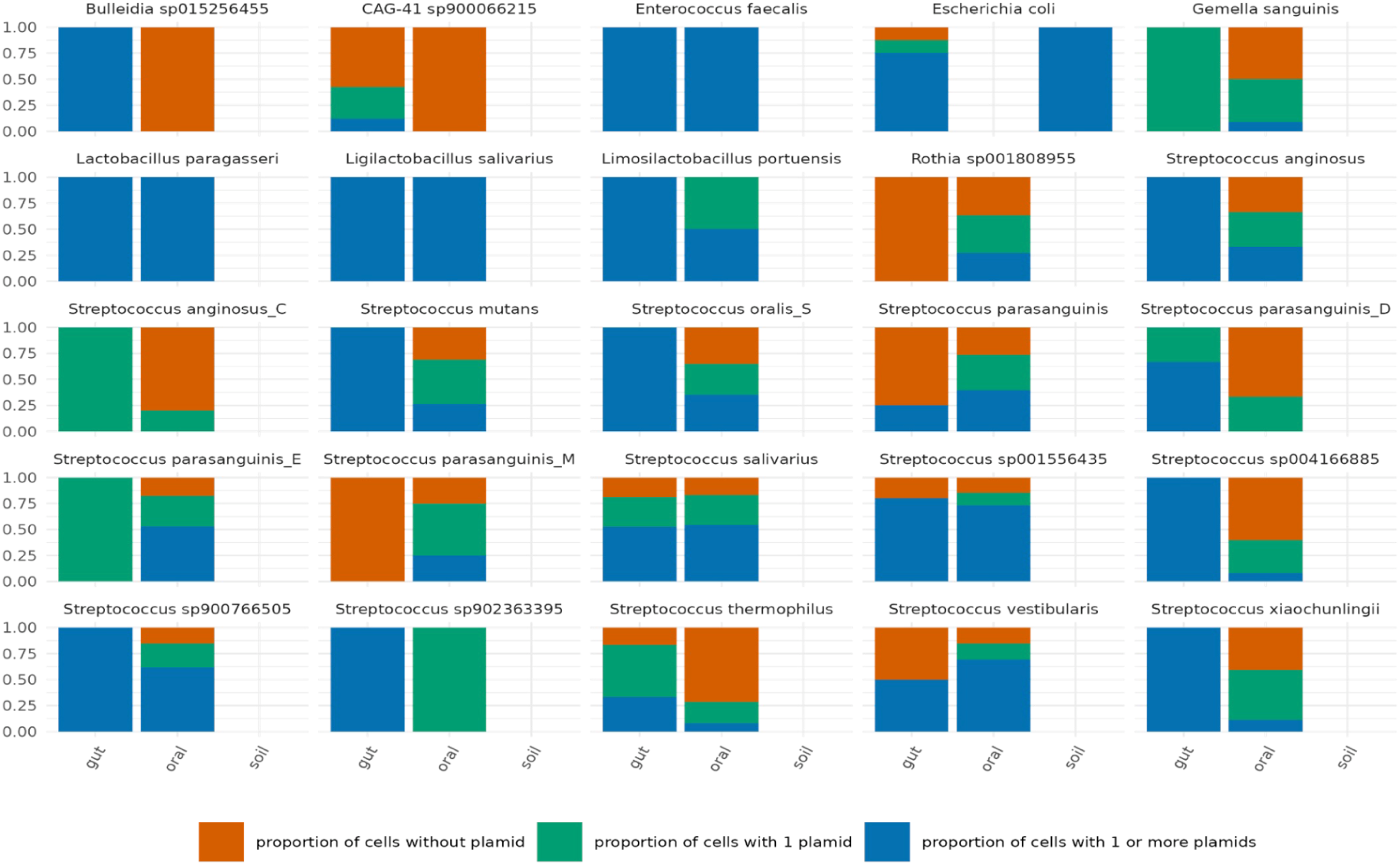
Prevalence of plasmid clusters shared among identical host species across diverse ecosystems.

**Supplementary Fig. 7.**
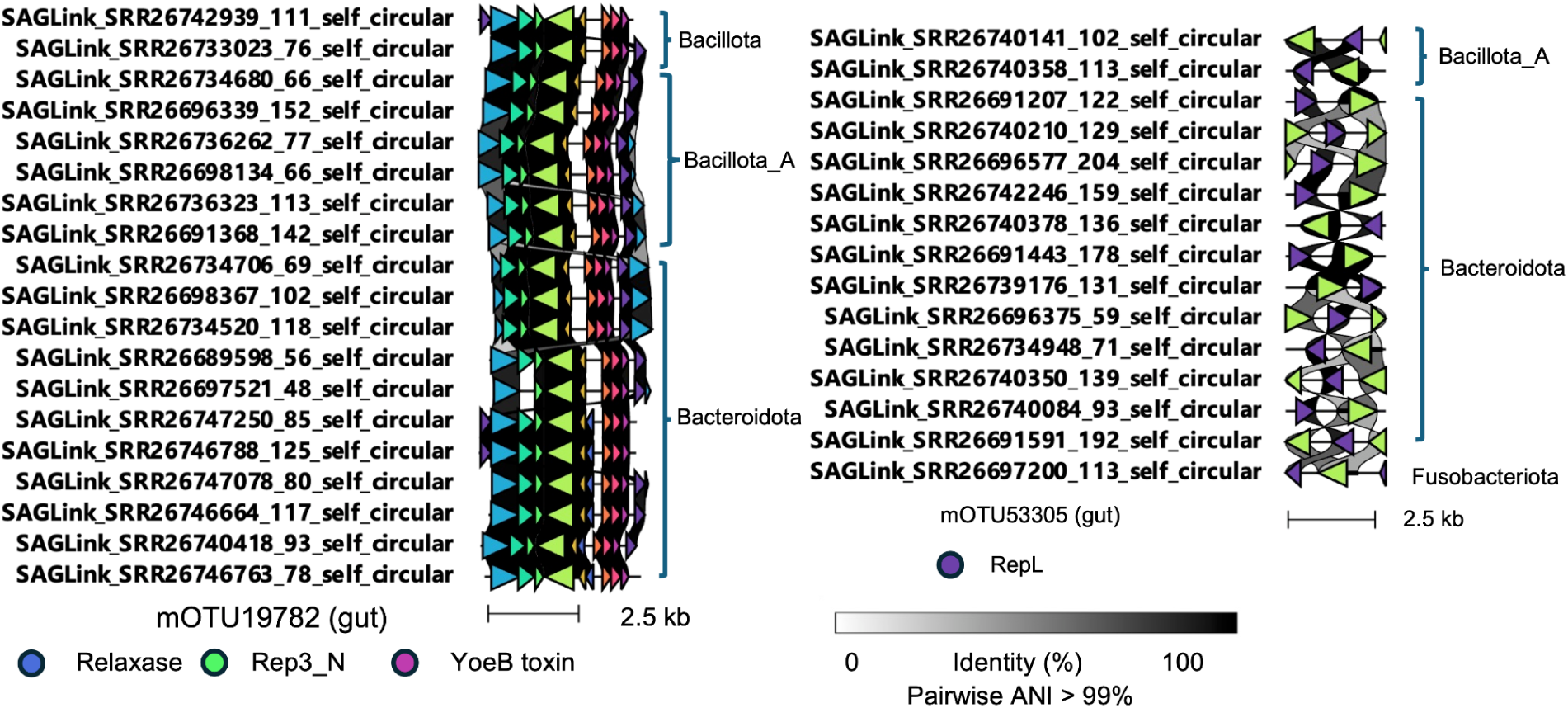
Additional circular and complete MGE sequences with host ranges spanning multiple phyla, identified beyond those shown in. Fig. 2d.

**Supplementary Fig. 8.**
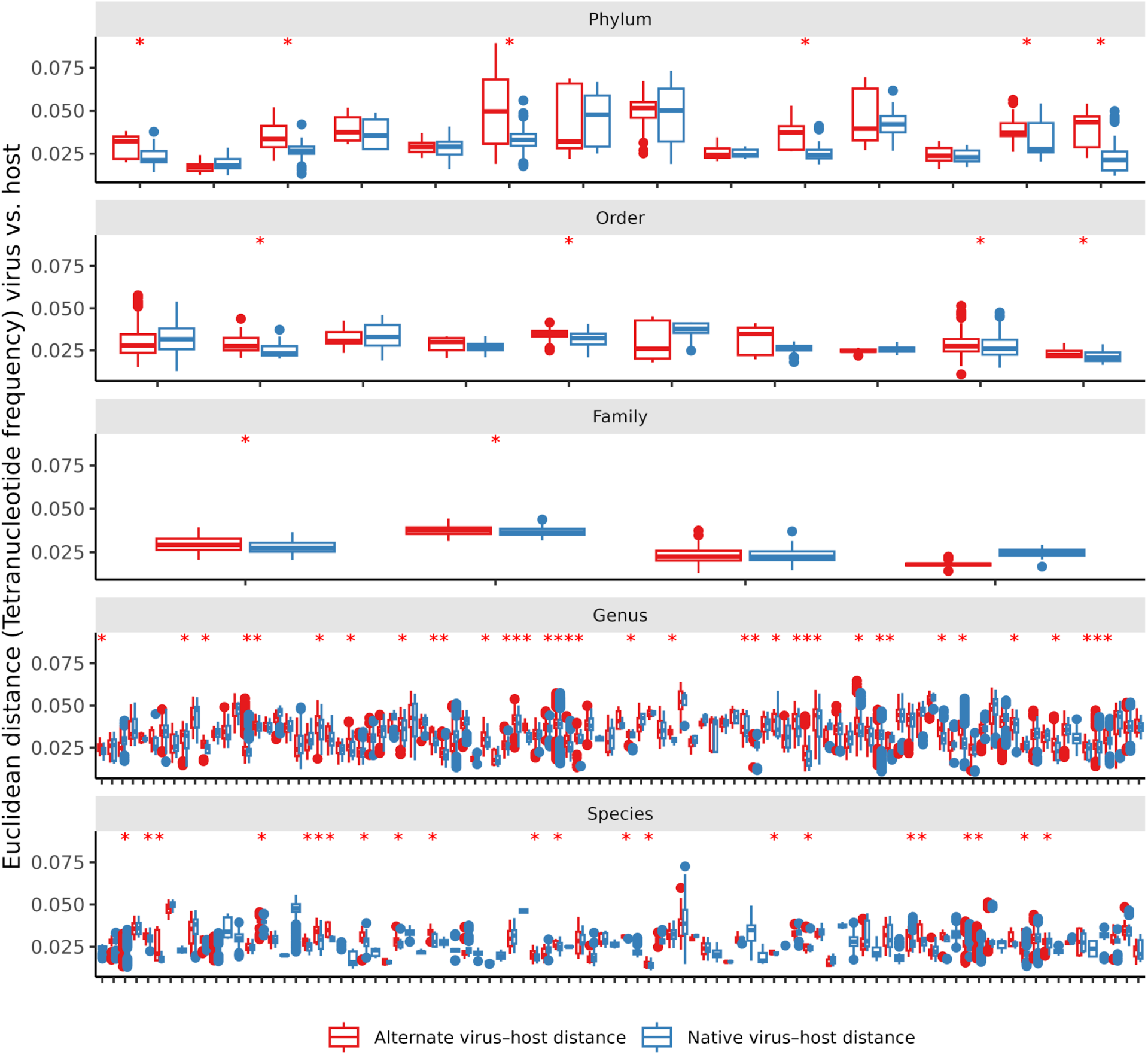
Comparison of primary and alternative virus–host tetranucleotide frequency distances across broad host-range vOTUs. For each vOTU with evidence of broad host range, Euclidean distances between tetranucleotide frequency (TNF) profiles of viral contigs and host SAGs were calculated for (i) the primary host associated with each virus and (ii) alternative hosts belonging to distinct taxonomic groups within the host range rank. Boxplots depict the distribution of TNF distances for primary and alternative host comparisons. We performed a one-sided Wilcoxon rank-sum test with the alternative hypothesis that alternative host TNF distances are greater than primary host distances.. Significant vOTUs (*p* < 0.05) are marked with red asterisks above the corresponding boxplots.

**Supplementary Fig. 9.**
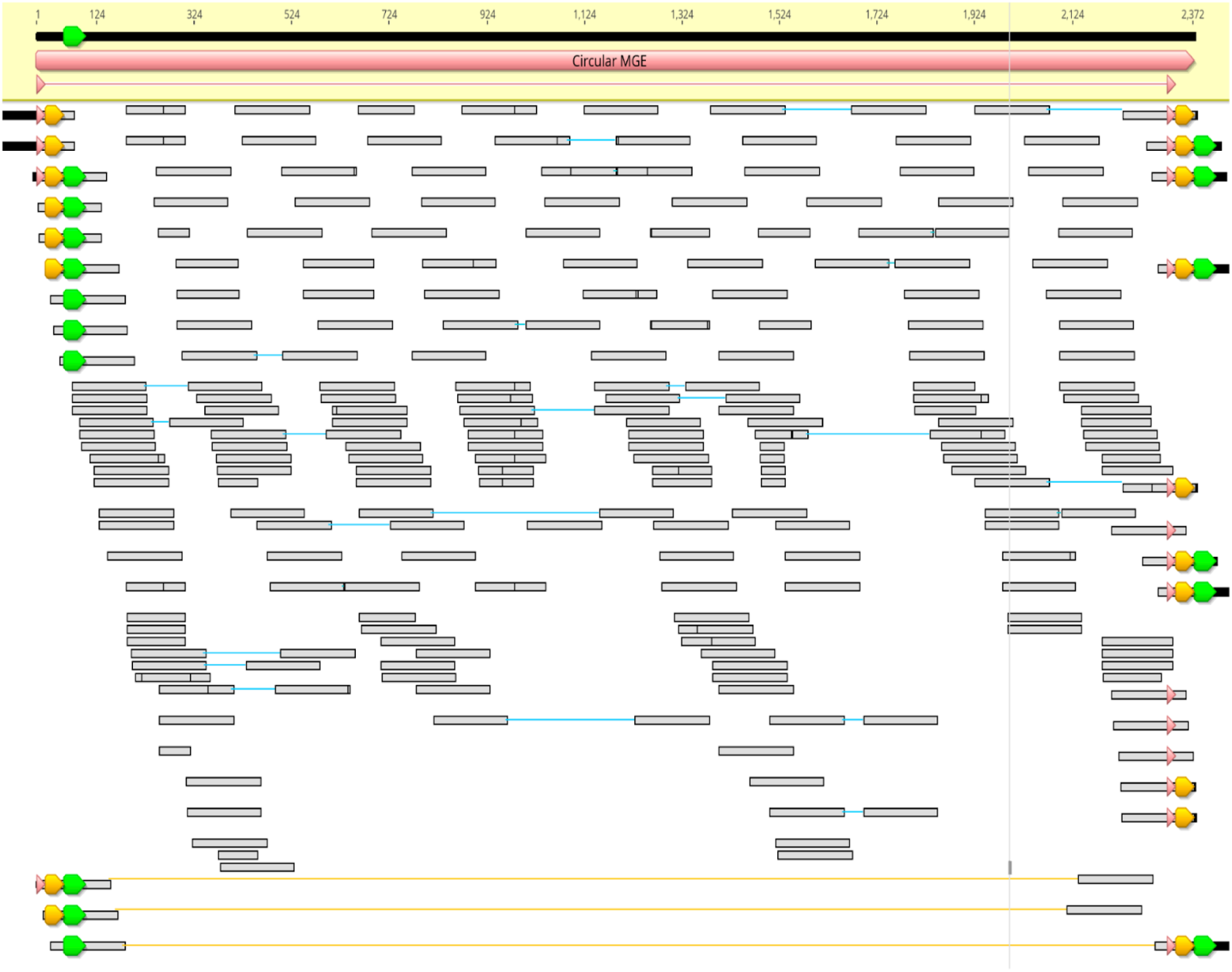
Manual curation of circular and complete non-canonical ECE encoding multidrug resistance proteins and exhibiting host ranges across multiple phyla, as shown in Fig. 4c. Arrows represent distinct nucleotide sequences, with identical colors indicating identical sequences. Vertical bars mark single-nucleotide polymorphisms (SNPs) relative to the reference genome. Reads connected by yellow lines span the sequence termini, providing evidence of circularity.

**Supplementary Fig. 10.**
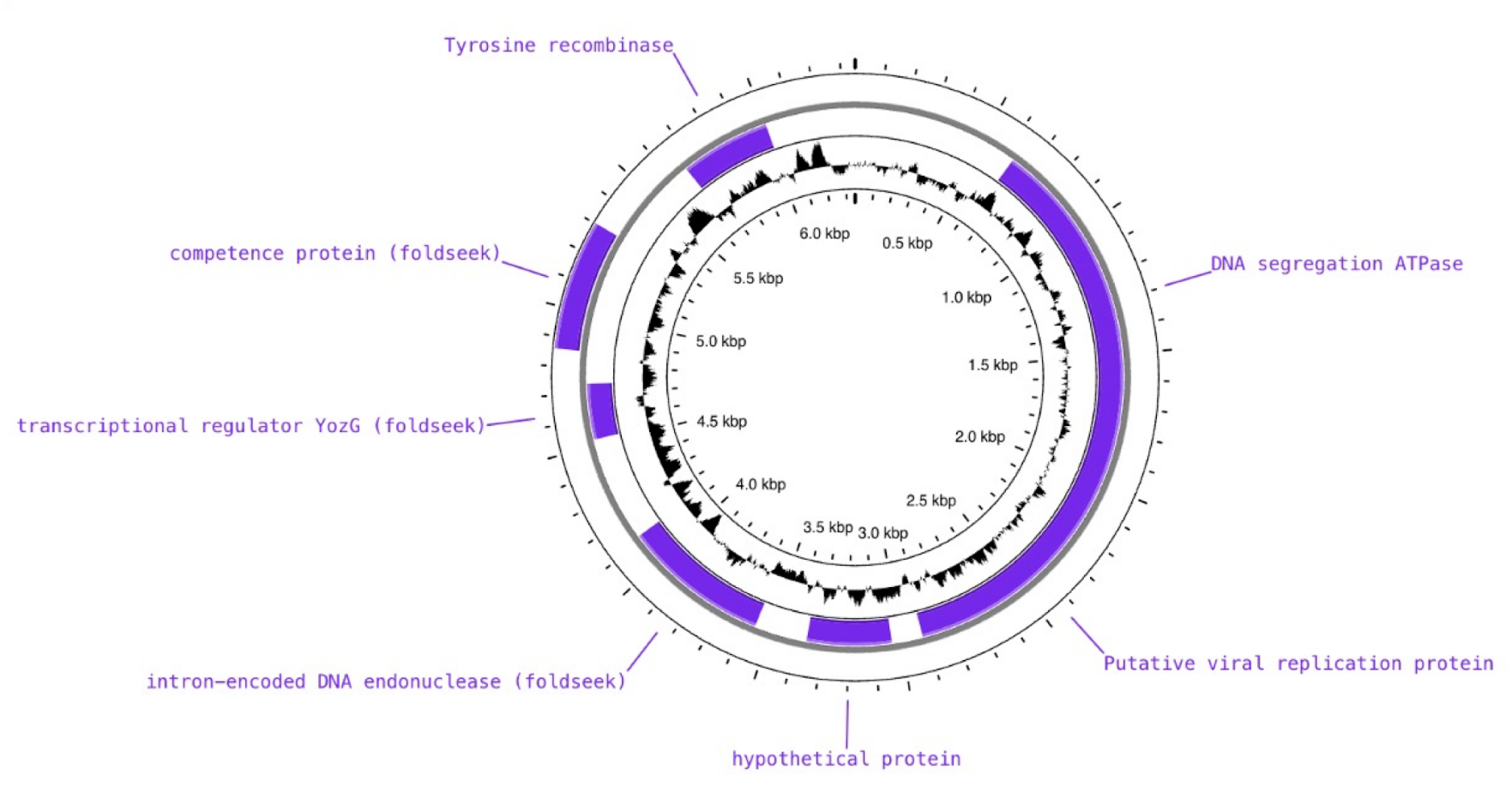
Non-canonical extrachromosomal elements (ECEs) with ecological significance identified from single-cell genome sequencing. ECEs targeted by CRISPR spacer sequences from 12 distinct SAGs of *Anaerostipes hadrus*, recovered across two independent sequencing projects.

**Supplementary Fig. 11.**
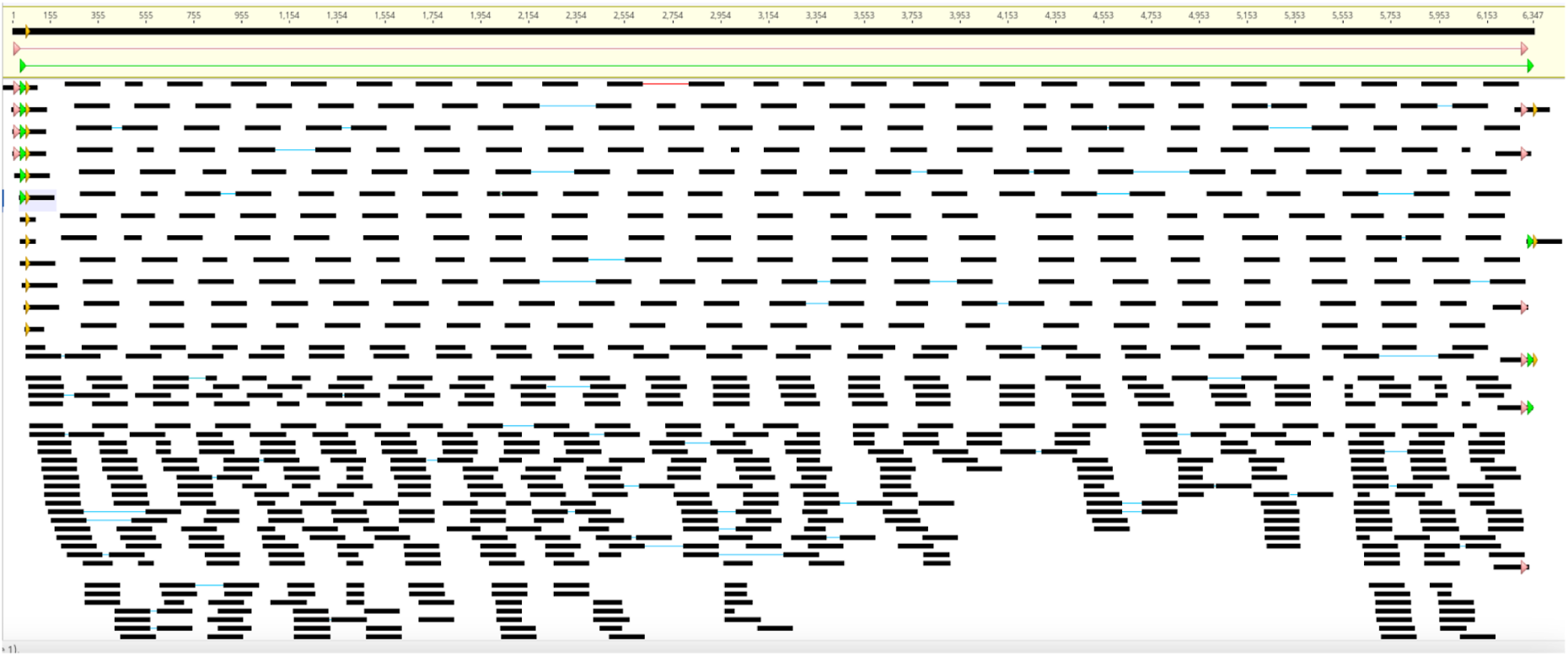
Manual curation of circular and complete non-canonical ECE targeted by multiple SAGs across two independent sequencing projects, as shown in Supplementary Fig. 10. Arrows represent distinct nucleotide sequences, with identical colors indicating identical sequences. Vertical bars mark single-nucleotide polymorphisms (SNPs) relative to the reference genome. Reads connected by yellow lines span the sequence termini, providing evidence of circularity.

**Supplementary Fig. 12.**
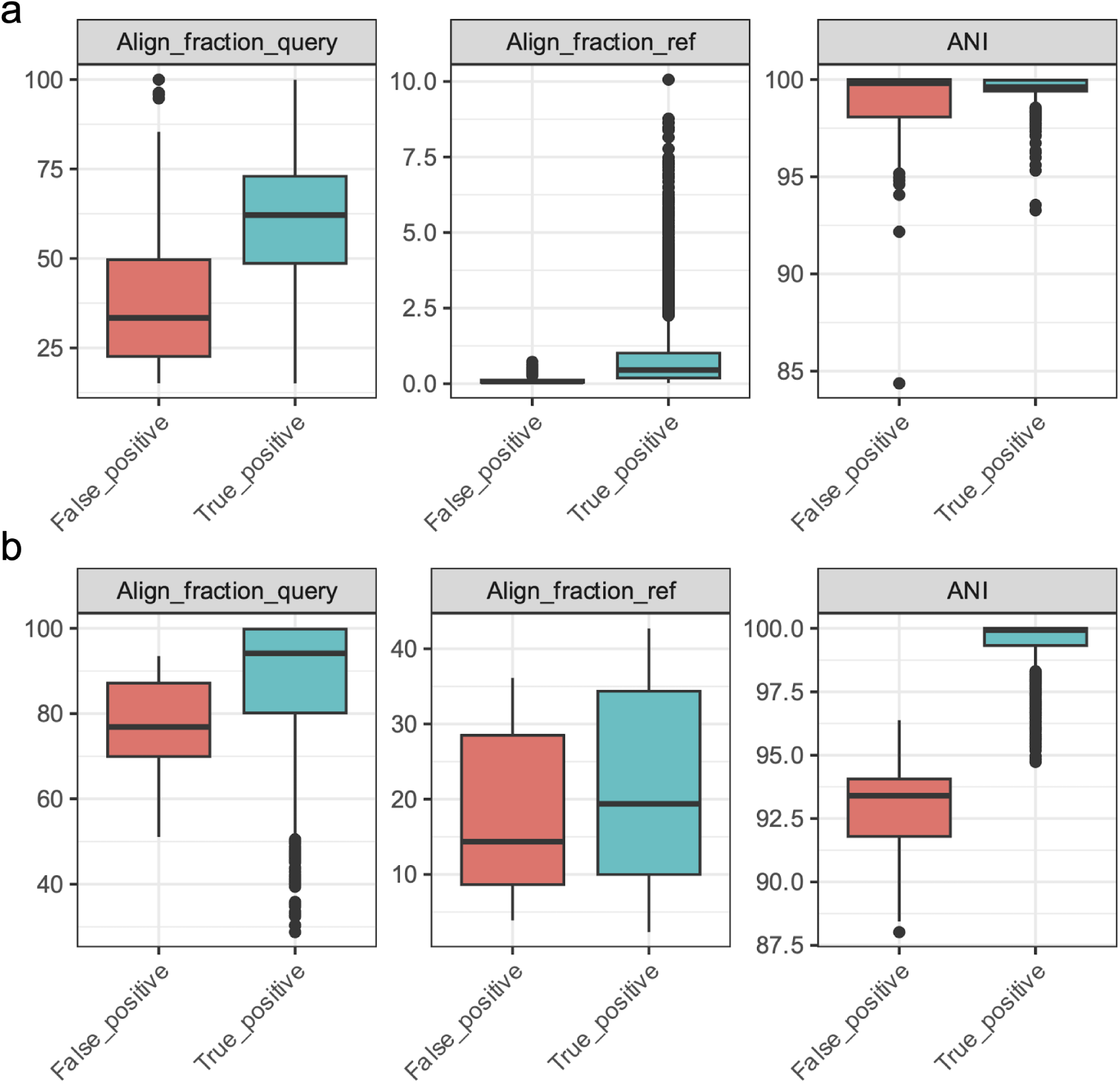
Benchmarking the Skani threshold used to determine SAG species identity. **a**, Aligned fraction and average nucleotide identity (ANI) between true and false positives among SAGs from a mock community generated in Zheng et al. (2022). **b**, Aligned fraction and ANI between true and false positives using simulated SAGs.

## Supplementary Tables

**Supplementary Table 1** Single cell genome sequencing data downloaded from NCBI

**Supplementary Table 2** Host assignment for the diverse host vOTUs and PTUs with host range across different species or above

**Supplementary Table 3** Annotations of the identified potential phage-plasmid genomes and their host rank

**Supplementary Table 4** Identified mobile gene elements with spacer hits

**Supplementary Table 5** Identified auxiliary genes in the MGEs

**Supplementary Table 6** Microbial strain with subpopulation-level MGE Variations

**Supplementary Table 7** Selected list of 529 species used to test the threshold for taxonomic assignment with skani.

